# Chemosynthetic Symbioses as Hidden Hubs of DMSP and Organosulfur Cycling in Marine Sediments

**DOI:** 10.64898/2025.12.05.692484

**Authors:** E. Kröber, K. Weinert, A. Mankowski, I.C. Özsefil, A. Porta Fidalgo, G. D’Angelo, C. Bannon, A.L. de Oliveira, H. Schäfer, N. Dubilier

## Abstract

Chemosynthetic symbioses between animals and bacteria are known to underpin productivity in the deep sea, yet the diversity of energy and carbon sources sustaining these associations in shallow-water environments remains poorly understood. Dimethylsulfoniopropionate (DMSP) is highly abundant in coastal habitats, where it is produced by seagrasses, phytoplankton, and heterotrophic bacteria, and occurs together with its breakdown product dimethyl sulfide (DMS) in shallow-water sediments. Here we show, supported by genomic and transcriptomic evidence, that DMSP and DMS cycling are integral to the energy and carbon metabolism of the gutless oligochaete *Olavius algarvensis* and its chemosynthetic symbionts. By assigning DMSP degradation pathways to individual members of the host’s microbial community, we reconstructed a network integrating demethylation and cleavage with energy conservation, methionine biosynthesis, and acetate assimilation into polyhydroxyalkanoates. We also identified a host-encoded methanethiol oxidase (MtoX) suggesting host participation in MeSH detoxification. Comparative metagenomic analyses of more than 60 gutless oligochaete species from globally distributed habitats showed that key DMSP- and DMS-processing genes (*dddP*, *dmdA*, *tmm*, *dmsA*) are widespread, indicating that organosulfur metabolism is a conserved feature of these symbioses. Our findings expand the recognized metabolic repertoire of shallow-water chemosynthetic symbioses and provide evidence that these associations directly contribute to marine DMSP and DMS cycling.

## Introduction

Organosulfur compounds, particularly dimethylsulfoniopropionate (DMSP) and its degradation product dimethyl sulfide (DMS), are central to marine sulfur cycling and climate regulation. DMSP is produced by phytoplankton, macroalgae, corals, seagrasses, salt-marsh plants, and diverse bacteria^1–5^. It serves multiple cellular functions, including osmoprotection, redox buffering, cryoprotection, herbivore deterrence, and carbon and sulfur storage^3,6–12^. Through enzymatic cleavage (Fig. 1), DMSP links the marine microbial loop to the atmosphere via DMS emissions, which account for up to 80% of global biogenic sulfur fluxes^13^, thereby influencing cloud formation, climate regulation, and even animal behavior^14–17^.

**Figure 1.**
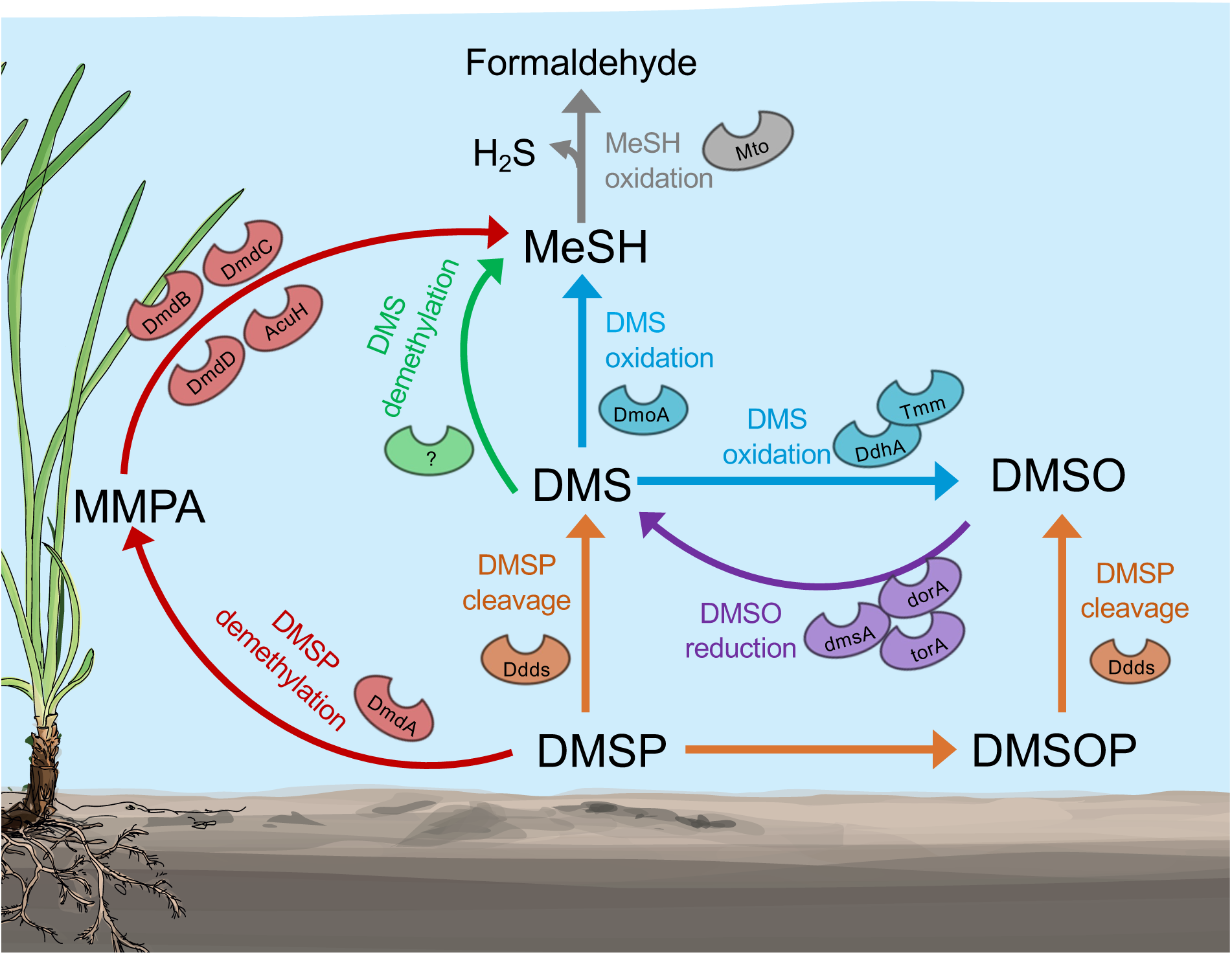
Major production and consumption pathways of OS compounds including known genes that encode key enzymes. DMSP demethylation (*dmdABCD*, *acuH*) generates methanethiol (MeSH). DMSP lyases (*ddd* genes) produce DMS from DMSP or DMSO from DMSOP. DMS is converted to MeSH via DMS monooxygenase (*dmoA*) or a yet unidentified DMS methyltransferase [40] (marked as “?”). DMS can also be oxidized to DMSO via DMS dehydrogenase (*ddhA*) or trimethylamine monooxygenase (*tmm*). DMSO can be reduced to DMS via DMSO reductases (*dmsA*, *dorA*) or via trimethyl amine N-oxide reductase (*torA*). MeSH can be degraded to H_2_S and formaldehyde by MeSH oxidase (*mtoX*). MMPA - 3-methylmercaptopropionate. Seagrass icon ©Elina Esken.

Despite intensive research on DMSP turnover in photic surface waters^13,18,19^, its role in shallow-water chemosynthetic symbioses remain largely unexplored. Many of these symbioses inhabit seagrass and coral reef sediments that are typically low in sulfide but rich in organosulfur and methylated amine compounds such as dimethylsulfoniopropionate (DMSP), dimethyl sulfide (DMS), and trimethylamine (TMA)^5,20–25^. These compounds could represent valuable alternative sources of carbon and energy in environments where conventional substrates for chemosynthesis, such as reduced sulfur compounds, are scarce.

Gutless oligochaetes are marine worms that lack digestive and excretory organs and rely entirely on chemosynthetic bacterial symbionts for nutrition and waste recycling^26–28^. Their symbionts exhibit high metabolic versatility, including sulfur oxidation, sulfate reduction, autotrophic C_1_ assimilation, and host metabolite recycling^27,29,30^. The most well studied species, *Olavius algarvensis*, hosts a consortium of up to five bacterial symbionts located beneath the cuticle, which are two sulfur-oxidizing Gammaproteobacteria (*Candidatus* Thiosymbion, “Gamma3”), up to three sulfate-reducing bacteria (Desulfobacteria^31^, referred to as “Delta1”, “Delta3”, or Desulfoconcordia^31^), and a spirochete^28,32^. The sulfur-oxidizing and sulfate-reducing symbionts form an internal sulfur cycle in which the thiotrophs oxidize reduced sulfur for carbon fixation while the deltaproteobacterial partners regenerate reduced sulfur compounds, maintaining tight redox coupling within the symbiosis.

Gutless oligochaetes thrive in oligotrophic, low-sulfide environments where access to alternative energy and carbon sources would provide a strong selective advantage. The abundance of organosulfur compounds such as DMSP and DMS in these habitats raises the possibility that they play a previously unrecognized role in sustaining these symbioses. Moreover, the vertical migration of gutless oligochaetes through the sediment column likely exposes them to DMSP-rich surface layers and suboxic zones enriched in DMS and its oxidation product dimethylsulfoxide (DMSO), creating dynamic opportunities for organosulfur cycling within this chemosynthetic symbiosis. While DMSP metabolism has been reported in heterotrophic symbioses, such as alphaproteobacterial scallop^33^ and gammaproteobacterial sponge symbionts^34^, it has not yet been demonstrated in chemosynthetic symbioses. This gap limits our understanding of how organosulfur compounds contribute to metabolic interactions and energy conservation within these systems.

Here we show that DMSP and DMS cycling are integral components of the symbiotic metabolism in the gutless oligochaete *O. algarvensis*. Using genome-resolved analyses and transcriptomics, we assigned DMSP degradation pathways to individual symbionts and reconstructed a network that links DMSP demethylation and cleavage to energy conservation, carbon assimilation and methionine biosynthesis. Building on this model system, we extended our analysis to over 60 gutless oligochaete species using comparative metagenomics, phylogenetic reconstruction, and gene-neighbourhood analyses to assess the diversity and distribution of DMSP- and DMS-processing genes. Together, these findings reveal that organosulfur metabolism is a conserved and ecologically relevant feature of shallow-water chemosynthetic symbioses.

## Material and Methods

### Symbiont-Resolved Screening in Olavius algarvensis

We performed a detailed, symbiont-resolved analysis of the consortium of *Olavius algarvensis* to identify the contributions of each bacterial partner to organosulfur cycling. Metagenome-assembled genomes (MAGs) from *O. algarvensis* metagenomes^35^ were screened for marker genes involved in the cycling of DMSP and DMS, as well as related intermediates (Fig. 1). Query genes (Supplementary Table S1) included those mediating DMSP cleavage (*ddd* family), demethylation (*dmdAB*), oxidation (*dmoA*, *ddhA*, *tmm*), DMSO reduction (*dmsA*, *dorA*/*torA*), and MeSH oxidation (*mtoX*). Searches were conducted using the BLAST software^36,37^. Candidate genes were manually verified through BLASTX searches against the NCBI nr and UniProt databases to confirm functional identity and remove spurious hits. Gene-specific thresholds for sequence length, identity, and e-value were applied (Supplementary Table S2), and verified sequences were subsequently used for phylogenetic reconstruction.

### Protein Structure Prediction and Comparative Analysis

The tertiary structures of MtoX homologs from *Olavius algarvensis*, *Homo sapiens* (SELENBP1), Caenorhabditis elegans (semo-1), and Methylophaga thiooxidans were predicted using AlphaFold2^38^ (v2.3.2) with the monomer model pre-set. For each sequence, the standard AlphaFold pipeline was applied, including iterative database searches with JackHMMER^39^ (UniRef90, MGnify) and HHblits^40^ (BFD, UniRef30) to generate multiple sequence alignments (MSAs), and HHsearch against the PDB70 database to identify structural templates. Five models were generated per sequence, and prediction confidence was assessed using Predicted Local Distance Difference Test (pLDDT) scores, which estimate residue-level positional accuracy. Protein sequences were aligned using Clustal Omega^41^ (Geneious Prime, https://www.geneious.com) to identify conserved motifs and catalytic residues characteristic of the MtoX/SELENBP1 family. Residue conservation was verified against the predicted tertiary structures. Structural visualization and comparison were performed in PyMOL^42^ (v2.5.5). Pairwise Cα superpositions were conducted using the *O. algarvensis* MtoX homolog as the reference to calculate Root-Mean-Square Deviation (RMSD) values, allowing quantitative assessment of fold conservation across homologs. Predicted structures were compared to evaluate whether the *O. algarvensis* MtoX homolog retains the canonical MtoX/SELENBP1 fold, thereby assessing its structural plausibility as a functional methanethiol oxidase.

### Transcriptomics

Specimens of *Olavius algarvensis* were collected near Elba (Sant’Andrea), Italy, in 2023. For each of three libraries, RNA was extracted from three pooled individuals. RNA extraction, rRNA depletion, and library preparation were performed by Novogene GmbH, with host rRNA removed to enhance the recovery of symbiont transcripts.

Adapter sequences from paired-end reads were trimmed using BBDuk^43^ (BBTools v39.08) with parameters *ktrim=rl mink=11 hdist=1 k=23 tbe tbo*, and read quality was assessed with FastQC^44^ (v0.12.1). Cleaned reads were mapped to a custom database containing the coding sequences of the six *O. algarvensis* symbionts using Bowtie2^45^ (v2.5.1) with the settings *-p 40 --no-unal -no-mixed --no-discordant -k 1 --un-conc-gz*. Resulting alignments were converted to BAM format with SAMtools^46^ (v1.21).

Transcript abundances were quantified as transcripts per million (TPM) using Salmon^47^ (v1.10.3). TPM values were imported into R via the tximport package (v1.37.2).

To identify and quantify genes involved in organosulfur cycling, a custom protein reference set was compiled from UniProtKB, and a tBLASTn search (v2.14.1+) was conducted against the symbiont coding sequence database with an e-value cutoff of 1e-40. For comparison, protein references of selected metabolic and housekeeping genes were retrieved from UniProtKB and analyzed with the same parameters. Hits with e-values < 1e-80 were further verified by BLASTx searches against the NCBI clustered NR database. Verified genes were screened in the expression dataset within R, and TPM values > 0 were visualized using ggplot2^48^ (v4.0.0).

### Gene mining across gutless oligochaetes

To assess the broader distribution of organosulfur cycling genes, we conducted a comprehensive survey of 207 metagenomes from over 60 gutless-oligochaete host species^35^. The same curated BLAST pipeline applied in *O. algarvensis* was used to identify marker genes associated with DMSP and DMS cycling (Supplementary Table S1). Candidate genes were validated as described above in „*Symbiont-Resolved Screening in* Olavius algarvensis”

### Symbiont-level gene distribution and enrichment analysis

To assess how organosulfur genes are distributed among symbiont lineages, contigs containing DMSP-, DMS-, or MeSH-cycling genes were searched against a reference database of metagenome-assembled genomes (MAGs) from 60 gutless oligochaete metagenomes^35^. Contigs not assigned to a MAG were categorized as “unbinned.” We counted the number of gene hits per symbiont lineage and visualized their relative abundance in a bubble plot. To test whether particular symbionts were statistically enriched in specific genes, we performed a chi-squared test on the contingency table of symbionts × genes and calculated Z-scores from the standardized residuals. Positive Z-scores indicate enrichment, while negative values indicate depletion.

### Phylogenetic and gene-neighbourhood analysis

Curated sequences were aligned in Geneious Prime v2021.0.3 using Clustal Omega (Sievers & Higgins, 2011/2018) and exported in PHYLIP format. Maximum-likelihood phylogenies were inferred with IQ-TREE^49^ (v1.6.12), using ModelFinder^50^ for substitution-model selection and ultrafast bootstrap for branch support (UFBoot^51^). Phylogenetic trees were visualized and annotated using iTOL^52^.

For gene-neighbourhood analysis, we examined up to 20 kb around each curated hit to identify co-localized genes potentially involved in sulfur, C_1_, acrylate, or redox metabolism. Genomic contexts were extracted and compared across symbionts and references to assess potential functional linkages and operon structures.

## Results and Discussion

### DMSP degradation and organosulfur cycling in Olavius algarvensis symbiosis

To understand organosulfur cycling (Fig. 1) in gutless oligochaetes, we first investigated the well-studied *Olavius algarvensis* symbiosis. This species harbours a stable and consistent symbiont consortium, enabling us to link specific organosulfur metabolic pathways to individual partners and reconstruct how each contributes to the turnover of DMSP, DMS, and related sulfur compounds within the holobiont.

Metagenomic analyses revealed that several of these symbionts encode key genes for DMSP degradation, DMS oxidation, and DMSO reduction, together forming a syntrophic cascade of organosulfur transformations within the consortium (Fig. 2a).

**Figure 2.**
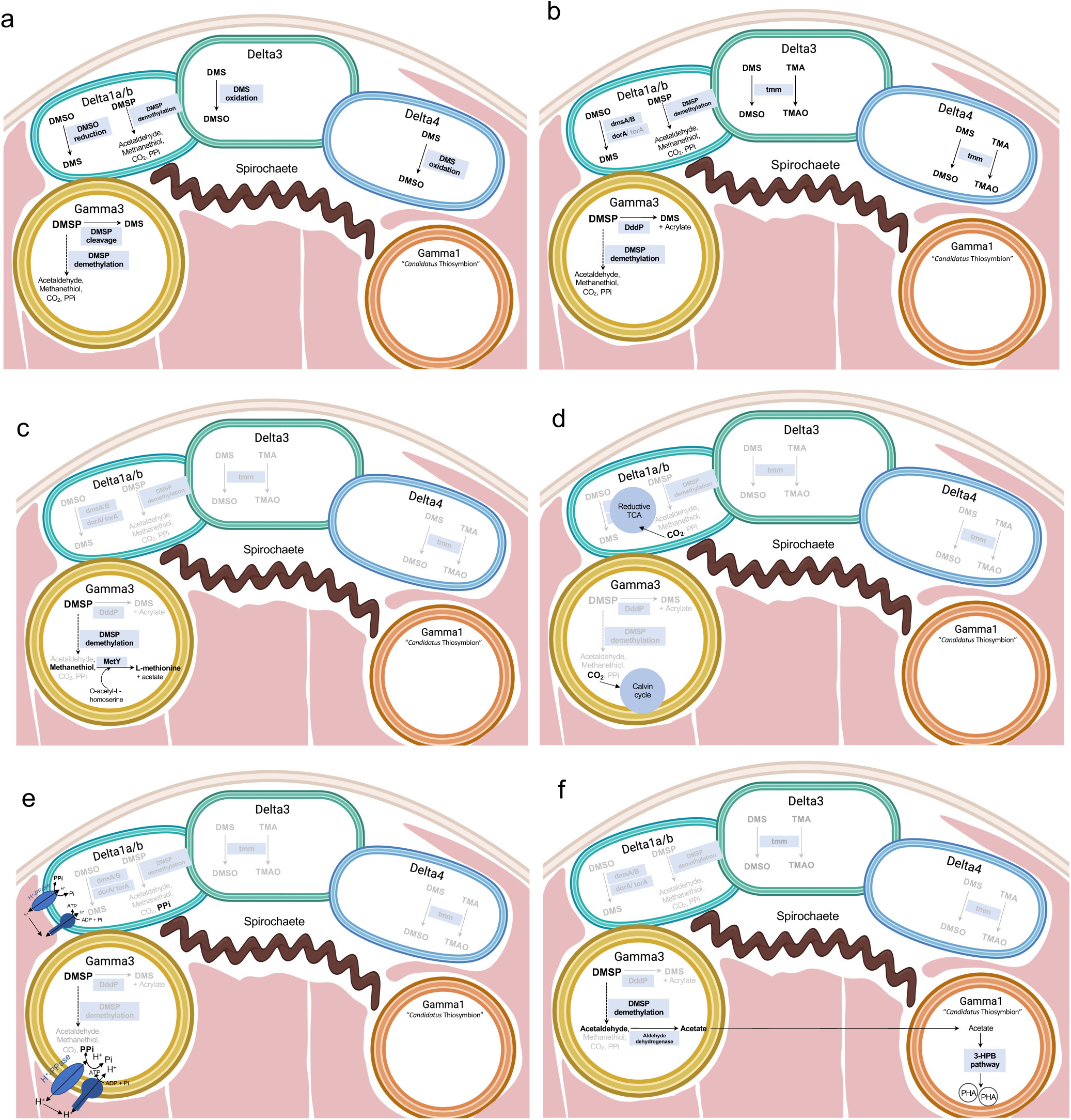
Overview of symbiotic DMSP/DMSO/DMS and MeSH metabolism based on metagenomics and metatranscriptomics in *O. algarvensis*. **(a)** Overview of the metabolic capability to cycle the OS compounds DMSP, DMSO, DMS in the sulfate-reducing (Delta1a/b, Delta3, Desulfoconcordia) and sulfur-oxidizing (*Ca.* Thiosymbion and Gamma3) symbionts. **(b)** Overview of the genetic capacity to cycle OS compounds in *Olavius algarvensis*’ symbionts. **(c)** MeSH produced during DMSP demethylation can be used for L-methionine production in the Gamma3 symbiont. **(d)** CO_2_ generated via the degradation of DMSP could be fixed into organic carbon by the autotrophic symbionts. **(e)** Pyrophosphate, also generated via the degradation of DMSP, could drive ATP production in Delta1 and Gamma3 symbionts. **(f)** Acetaldehyde from DMSP degradation could lead to acetate production in Gamma3 symbionts, which in turn could drive polyhydroxyalkanoate (PHA, bioplastic) production in the *Ca*. Thiosymbion symbiont. Microbial icons were created with BioRender.

Within this cascade, the Gamma3 and Delta1 symbionts play central roles, possessing the potential to demethylate DMSP via the *dmdABC* and *acuH* pathway, yielding acetaldehyde, CO_2_, pyrophosphate (PPi), and MeSH (Fig. 2a,b). Notably, the Gamma3 symbiont encodes genes for both the DMSP demethylation (*dmdABC*, *acuH*) and cleavage (*dddP*) pathways, indicating metabolic flexibility in DMSP degradation (Fig. 2a,b). The Gamma3 symbiont further encodes *metY*, enabling the conversion of MeSH and O-acetylhomoserine into L-methionine, thus providing a sulfate-independent route for methionine biosynthesis (Fig. 2c).

The CO_2_ released during DMSP degradation could be assimilated via autotrophic carbon fixation pathways. Gamma3 encodes the Calvin-Benson-Bassham cycle^27^, whereas Delta1 encodes the Wood-Ljungdahl (reductive acetyl-CoA) pathway^53^, a key CO_2_ fixation route typical of sulfate-reducing bacteria. This suggests that DMSP-derived carbon may be incorporated into biomass via distinct fixation mechanisms in sulfur-oxidizing (Gamma3) and sulfate-reducing (Delta1) symbionts (Fig. 2d). Pyrophosphate (PPi), another demethylation by-product, may fuel ATP synthesis via H⁺-translocating pyrophosphatases (H⁺-PPases) encoded by both Delta1 and Gamma3 symbionts^27^, enhancing energy conservation (Fig. 2e). Acetaldehyde, another intermediate, can be oxidized to acetate, which may enter the 3-hydroxypropionate bi-cycle (3HPB) of the Gamma1 symbiont and contribute to polyhydroxyalkanoates (PHA) storage, a major carbon reserve in this lineage (Fig. 2f).

The flavin-dependent monooxygenase Tmm, which oxidizes both trimethylamine (TMA) to trimethylamine-N-oxide (TMAO) and DMS to DMSO, is encoded by the Delta3 and Desulfoconcordia symbionts (Fig. 2b). Given that both DMS and TMA are abundant in coastal sediments and can serve as reduced methylated substrates, this enzyme likely enables these symbionts to participate in methylated-amine and sulfur oxidation within the consortium. The cooccurrence of *tmm* with DMSO-reductase genes (*dmsABC*) supports the potential for a DMSDMSO redox loop under fluctuating sediment redox conditions.

The fate of MeSH, produced either by symbiotic DMSP demethylation or from external sediment sources where it is generated through microbial DMS and methionine catabolism, is less clear. While this reduced sulfur compound could theoretically be oxidized to H_2_S, no *mtoX* homolog was found in any symbiont. However, a candidate *mtoX* gene was identified in the host genome (Supplementary Fig. S1, S12a), suggesting that MeSH detoxification may be carried out by the animal itself. Phylogenetic analysis placed the *O. algarvensis* homolog in a sister clade to bacterial MtoX, clustering with verified animal SELENBP1 sequences (Fig. S12a). Sequence alignment (Fig. S1a) revealed conservation of two catalytic residues (G272 and H380) essential for copper coordination and MeSH oxidation^54,55^. AlphaFold2 structural predictions (Fig. S1c,d) showed that the *O. algarvensis* homolog adopts the canonical eight-bladed β-propeller fold characteristic of the MtoX/SELENBP1 family, indicating that it retains the overall architecture. Host-derived *mtoX* transcripts were detected in epidermal cells (data not shown), consistent with the localization of symbionts directly beneath the cuticle in the inter-epidermal region, suggesting a potential spatial coupling between symbiont MeSH release and host detoxification. The potential host-mediated oxidation of MeSH in *O. algarvensis* parallels processes in mammals, where MeSH produced by gut microbiota or endogenous metabolism is oxidized by the host enzyme methanethiol oxidase (SELENBP1)^54^. Host-mediated MeSH oxidation could thus represent metabolic complementarity, in which symbionts produce MeSH and the host detoxifies it, potentially regenerating sulfane sulfur or H_2_S that could fuel sulfur-oxidizing symbionts. While currently hypothetical, this mechanism warrants experimental validation.

Altogether, organosulfur metabolism in *O. algarvensis* is both taxonomically partitioned and metabolically integrated. DMSP acts as a central substrate fuelling methionine production, carbon fixation, acetate assimilation, PHA biosynthesis, and sulfur redox cycling. The possible host contribution to MeSH oxidation underscores that organosulfur transformations likely span both symbiont and host. These findings establish a model in which DMSP and DMS metabolism add a new layer to the canonical sulfur cycling between sulfur oxidizers and sulfate reducers. In this model, DMSP-derived compounds provide flexible carbon and energy sources, while DMS and DMSO function as reversible redox intermediates, buffering the consortium under fluctuating sediment oxygen levels.

### Organosulfur pathways are transcriptionally active across *O. algarvensis* symbionts

To assess whether the organosulfur pathways identified in *O. algarvensis* metagenomes are actively expressed under natural conditions (field-fixed samples), we analysed symbiont-derived transcriptomes. Key genes involved in DMSP degradation, DMS/DMSO interconversion, and methionine biosynthesis, including *dddP*, *dmdA*, *dmdB*, *dmdC*, *acuH*, *dmsA*, *dmsB*, and *metY*, were transcribed in both the Gamma3 and Delta1 symbionts (Fig. 3a), confirming their functional relevance *in vivo*.

**Figure 3.**
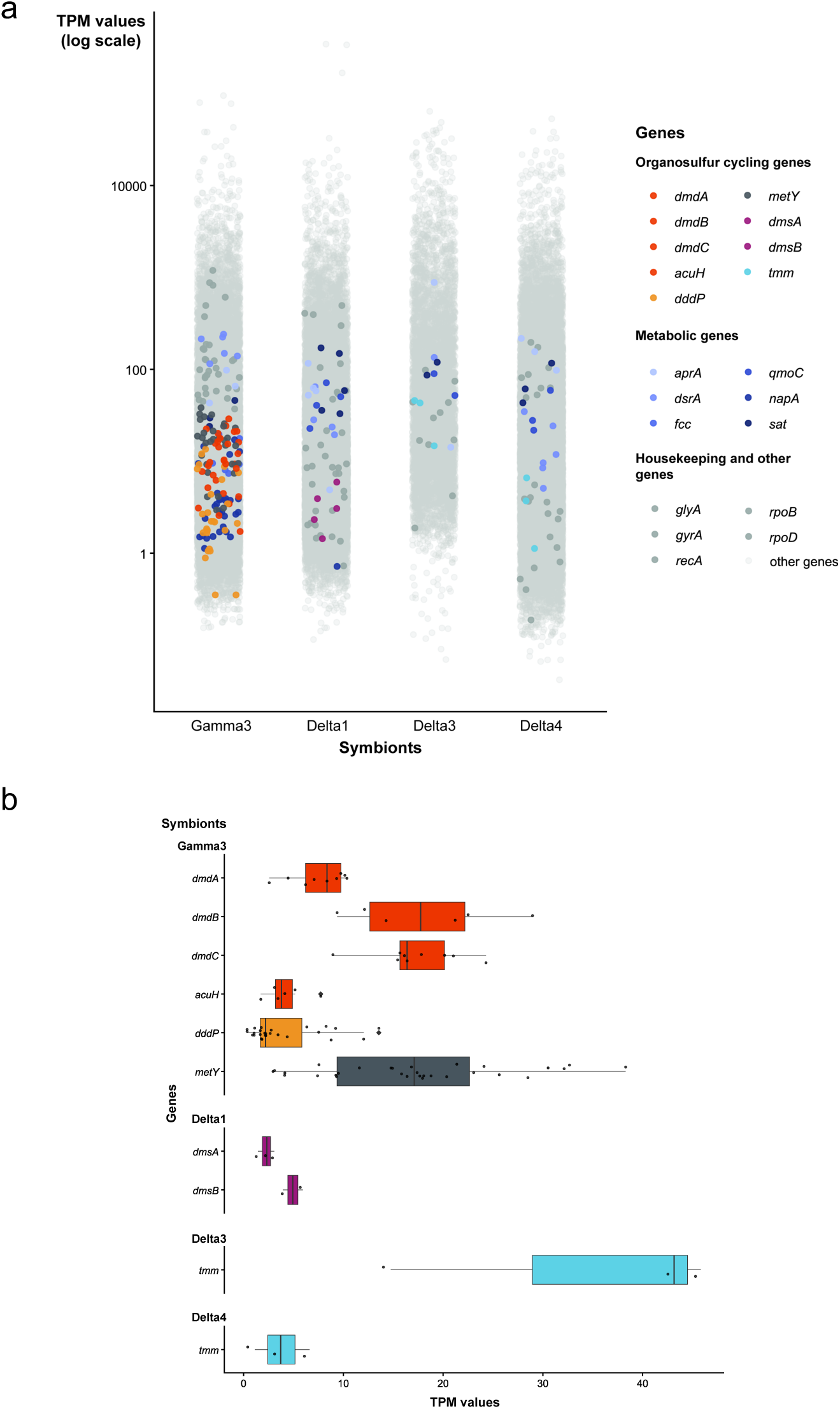
Expression of organosulfur metabolism genes in symbionts of *Olavius algarvensis*. **(a)** Transcript abundance (TPM, log scale) of all genes encoded by the different symbionts of *O. algarvensis*. Genes involved in organosulfur metabolism are highlighted in color, including *dddP*, *dmdA*, *dmdB*, *dmdC*, *acuH*, *dmsA*, *dmsB*, and *metY*. Housekeeping genes and general metabolic genes are indicated in grey tones. **(b)** Expression levels (TPM) of selected organosulfur metabolism genes across four symbionts: Gamma3, Delta1, Delta3, and Desulfoconcordia. *Ca.* Thiosymbion and Spirochaeta symbionts are not shown as they lacked detectable expression of organosulfur genes in *O. algarvensis*. While gene expression levels are generally low, several pathways are consistently transcribed across symbionts. Notably, *tmm* expression in Delta3 is the highest among all surveyed genes.

A broader comparison across all organosulfur-competent symbionts in *O. algarvensis* (Fig. 3b) revealed that expression levels were generally lower than those of core sulfur oxidation and reduction pathways (e.g., *aprA*, *dsrA*), but within the range of several housekeeping (*glyA*, *gyrA*, *recA*, *rpoB*) or nitrate reduction genes (e.g., *napA*), and consistent across biological replicates. Among all surveyed organosulfur genes, *tmm* expression in the Delta3 symbiont was highest, suggesting a particularly active role in DMS oxidation and/or methylated amine metabolism. Although absolute transcript abundances were low, the repeated recovery of these transcripts across replicates indicates stable and recurrent pathway activity. Such constitutive but low-level expression may represent a metabolic readiness strategy, enabling symbionts to rapidly respond to transient DMSP or DMS inputs, a pattern reminiscent of proteorhodopsin expression in SAR11 bacteria, which maintains preparedness under resource-fluctuating conditions^56^. It is important to note that these transcriptomes were from pooled *Olavius* individuals, and thus represent population-level averages rather than single-symbiont expression profiles. Nevertheless, the consistent detection of organosulfur transcripts across independent samples underscores their ecological and physiological importance. Together, these results demonstrate that organosulfur cycling in *O. algarvensis* is not only genetically encoded but transcriptionally active, forming a core metabolic framework that supports energy conservation and redox balance within the symbiosis.

### Organosulfur compound cycling genes are widespread in metagenomes of oligochaete symbionts

To assess whether this organosulfur-based metabolic integration is unique to *O. algarvensis* or a broader feature of gutless oligochaete symbioses, we expanded our analysis across more than 60 gutless oligochaete species from globally distributed habitats, screening 269 metagenomes for homologs of key pathway genes (Fig. 1,4). Because gutless oligochaetes host diverse combinations of secondary symbionts that vary across species and environments, differences in gene presence likely reflect lineage-specific metabolic repertoires rather than pathway loss. This analysis revealed pronounced contrasts in gene representation. Some genes, including *dddP*, *dmdA*, *dmdB*, and *tmm*, were widely distributed, detected in 46, 37, 39 and 37 host species, respectively, and often present as multiple homologs per individual. *dmsA* was also common (22 species), though less evenly represented. In contrast, *mtoX* showed a more restricted distribution, appearing slightly more prevalent in (but not limited to) the *Inanidrilus* clade of the host phylogeny (Fig. 4). By contrast, several lyase and oxidation markers (*dddD*, *dddL*, *dddY*, *dddQ*, *dddW*, *dmoA*, and *ddhA*) were rare or absent, detected in only a few cases (4, 0, 0, 0, 0, 1 and 1 species, respectively). Together, these patterns highlight a core set of widely conserved DMSP- and DMS-processing genes, contrasted by a small number of sporadic or lineagespecific enzymes. This suggests a modular architecture of organosulfur metabolism, in which some reactions are broadly maintained while auxiliary pathways vary among symbiont communities.

**Figure 4.**
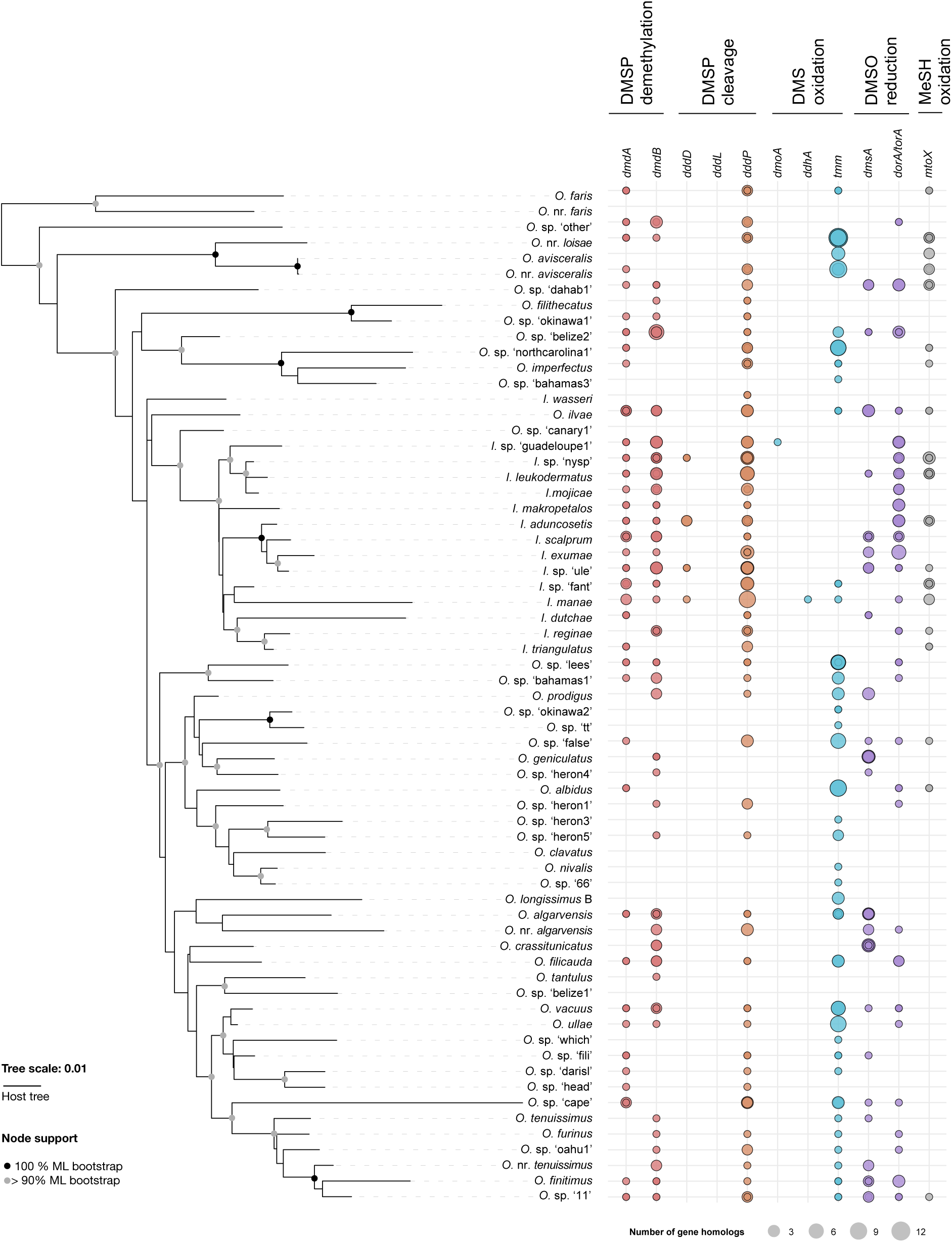
Genes for OS compound cycling are widely distributed across gutless oligochaetes. 207 gutless oligochaete metagenomes from *Olavius* and *Inanidrillus* species were analyzed for the presence of genes involved in OS cycling (for overview see Figure 1). On the left side a maximum-likelihood tree of the host 28S rRNA gene phylogeny is shown for the gutless oligochaete species. Nodes with none-parametric bootstrap support > 90% are highlighted in grey and black. The right panel shows the presence and abundance of homolog genes involved in OS cycling for each host species. Only manually curated hits were considered.

To examine how these genes are distributed among specific symbiont lineages, we mapped DMSP-, DMS-, and MeSH-cycling genes onto 60 gutless oligochaete metagenomes (Supplementary Fig. S11a). Given the compositional diversity of symbiont consortia across hosts, this approach allowed us to determine whether organosulfur gene distributions align with particular bacterial lineages rather than being randomly dispersed. Genes were assigned to symbionts based on their contig origin, revealing clear lineage-specific patterns. An enrichment analysis (Supplementary Fig. S2a) confirmed that Gamma3, several Deltaproteobacteria, and multiple Alphaproteobacteria lineages are significantly enriched in specific gene sets, consistent with their specialized metabolic roles within the consortium. These results demonstrate that organosulfur functions are non-randomly partitioned among symbiont lineages, reflecting metabolic complementarity and evolutionary adaptation to distinct ecological niches within the holobiont.

### DMSP cleavage and demethylation pathways are widespread among gutless oligochaete symbionts

Metagenomic analyses revealed that gutless oligochaete symbionts possess both major DMSP degradation routes, cleavage and demethylation, distributed across multiple bacterial partners (Fig. 4). The DMSP lyase *dddP* was particularly common and phylogenetically diverse, whereas *dddD* was detected only rarely. Gene neighbourhoods associated with *dddP* in symbiont contigs linked DMSP cleavage to redox and nitrogen metabolism, suggesting integration of DMS production with other central metabolic processes (Supplementary Results 1; Fig. S3-S4). The alternative demethylation pathway, encoded by *dmdA* and *dmdB*, was also prevalent (detected in 37 and 39 host species, respectively, Fig. 4) but displayed lineage-specific partitioning. Alphaproteobacterial symbionts typically encoded both genes,

Gammaproteobacteria mainly *dmdA*, and Deltaproteobacteria predominantly *dmdB*, indicating a division of labour in DMSP demethylation or differential gene retention or loss among symbiont groups (Supplementary Results 2; Fig. S5-S6). These patterns suggest complementary metabolic roles among symbionts and highlight DMSP catabolism as a key axis of interspecies cooperation within the consortium.

### DMS oxidation and DMSO reduction form a coupled redox system

Genes for DMS oxidation were recovered across most host species but differed strongly in prevalence. *tmm* (trimethylamine monooxygenase) was by far the most abundant and phylogenetically widespread marker, whereas *dmoA* (DMS monooxygenase) and *ddhA* (DMS dehydrogenase) were rare (Supplementary Results 3, Fig. 4). Given Tmm’s dual activity on DMS and trimethylamine, these data indicate that Tmm-mediated oxidation represents the dominant DMS-to-DMSO pathway in gutless symbionts, linking sulfur and nitrogen metabolism (Supplementary Results 3; Fig. S7-S9).

DMSO reduction genes were also common and segregated between symbiont lineages: Deltaproteobacteria typically carried the *dmsABC* operon, whereas Alphaproteobacteria encoded periplasmic *torA/dorA*-type reductases. Together with the widespread *tmm*, these genes support a DMS-DMSO redox loop within the holobiont that could buffer redox fluctuations and sustain electron flow under variable oxygen conditions (Supplementary Results 4; Fig. S10-S11).

### Methanethiol oxidation involves both symbionts and host

Beyond DMS and DMSO cycling, DMSP demethylation directly and DMSP cleavage indirectly (via DMS oxidation) generate methanethiol (MeSH), a potentially toxic intermediate. Genes encoding methanethiol oxidase (*mtoX*) were moderately common among symbionts and formed distinct alphaproteobacterial and gammaproteobacterial clades, each retaining putative copper-maturation systems (*mauG*, *SCO1*; Supplementary Results 5; Fig. S12). Additionally, the *O. algarvensis* host genome encodes a SELENBP1-like methanethiol oxidase with conserved catalytic residues, suggesting that both partners contribute to MeSH detoxification (Supplementary Results 5; Fig. S1).

### An integrated network of organosulfur metabolism

Taken together, gutless oligochaete symbionts encode a cohesive yet taxonomically partitioned network of organosulfur pathways linking DMSP degradation, DMS/DMSO interconversion, and methanethiol oxidation. These processes collectively provide redox flexibility and metabolic resilience in low-sulfide coastal sediments, positioning gutless oligochaete symbioses as previously unrecognized micro-scale hubs in marine organosulfur cycling.

## Conclusions

Our comparative analysis demonstrates that organosulfur metabolism genes are widespread yet unevenly represented across gutless oligochaete symbioses. Core genes such as *dddP*, *dmdA*, *tmm*, and *dmsA* are consistently present, whereas others (*dddD*, *dddL*, *ddhA*, *dmoA*) occur sporadically, reflecting lineage-specific specialization and modularity within these multi-partner consortia.

In *O. algarvensis*, we resolved distinct organosulfur pathways at the symbiont level, showing that DMSP and its degradation products are not minor intermediates but key substrates for energy and carbon metabolism, with genes actively transcribed in situ. Previous studies emphasized the internal cycling of reduced sulfur compounds such as hydrogen sulfide (H_2_S) between sulfate-reducing and sulfur-oxidizing symbionts. However, because ambient sulfide concentrations in the worms’ natural habitat are extremely low (< 1 µM)^32^, this loop alone cannot sustain the symbiosis. Our results therefore identify DMSP degradation as a complementary process that expands the metabolic foundation of the consortium.

Degradation through both the demethylation and cleavage pathways produces metabolites, including MeSH, CO_2_, pyrophosphate (PPi), acetaldehyde, and DMS, which could be exchanged among symbionts and integrated into core cellular processes. These include methionine biosynthesis, carbon fixation (via Calvin-Benson-Bassham and Wood-Ljungdahl pathway), ATP generation through H⁺-translocating pyrophosphatases, and acetate assimilation into polyhydroxyalkanoate (PHA) storage. The co-occurrence of both DMSP-degradation pathways in the Gamma3 symbiont underscores metabolic flexibility, paralleling that of *Pelagibacter ubique* (SAR11), which shifts between demethylation and cleavage depending on DMSP availability^57–59^. While bacterial symbionts appear to be the principal mediators of DMSP and DMS turnover, the detection of a host-encoded *mtoX*-like gene, which is also transcriptionally expressed in epidermal cells, suggests that the animal host may contribute to methanethiol oxidation. Such a cross-domain metabolic handoff could regenerate H_2_S or sulfane-sulfur for sulfur-oxidizing partners, illustrating metabolic coupling between host and symbionts, similar to host-microbe methanethiol oxidation in mammals^54^. Eukaryotic flavin-containing monooxygenases (FMOs), present in many metazoans including *Caenorhabditis elegans*, may provide additional, yet unexplored, routes for methylated amine or sulfur compound processing. These potential host contributions warrant targeted genomic, transcriptomic, and biochemical validation. Ecologically, gutless oligochaetes inhabit seagrass-associated sediments that are hotspots of DMSP production, yet their contribution to DMSP and DMS cycling has remained poorly characterized. Our findings indicate that these chemosynthetic symbioses harbour the genomic potential to transform volatile organosulfur compounds, linking host-associated microbial metabolism to localized sulfur and carbon transformations. The results suggest that gutless oligochaetes and their symbionts may influence organosulfur compound turnover and redox dynamics at the sediment-water interface, shaping micro-scale biogeochemical gradients rather than global elemental cycling. Although our evidence is primarily genomic and transcriptomic, it establishes a framework for experimental validation. In situ isotope tracing, enzyme assays, and spatial gene-expression analyses will be key to quantifying organosulfur fluxes within host tissues and surrounding sediments. Because chemosynthetic partnerships are widespread among marine animals, including lucinid clams, vestimentiferan tubeworms, bathymodiolin mussels, and sponges, it is plausible that DMSP and DMS cycling constitute a more general, yet previously underappreciated, feature of marine symbiotic metabolism. Altogether, our study reframes DMSP not merely as a marine osmolyte but as a central metabolite integrating sulfur, carbon, and energy flow in benthic chemosynthetic consortia, revealing a previously unrecognized biochemical link between microbial symbioses, sediment biogeochemistry, and the broader marine sulfur cycle.

## Supporting information

Supplementary Results

## Acknowledgements

This work was supported by the Emmy Noether Program of the DFG under project number 519818445, awarded to EK and the Max Planck Society.

We are thankful for sample collections and field assistance by Harald Gruber-Vodicka, Alexander Gruhl, Anna Ansebo, Anna Blazejak, Anne-Christin Kreutzmann, Christian Lott, Claudia Bergin, Dolma Michellod, Emilia M. Sogin, Erica Mejlon, Erich Mueller, Falk Warnecke, Fred Wells, Jörg Ott, Judith Zimmermann, Katrine Worsaae, Ken Halanych, K. B. Brandon Seah, Lisa Matamoros, Lena Gustavsson, Mario P. Schimak, Michael Hadfield, Miriam Sadowski, Miriam Weber, Olav Giere, Oliver Jäckle, Nicholas Bekkouche, Olivier Gros, Pamela Reid, Philippe Bouchet, Pierre De Wit, Ramon Rosello-Mora, Silke Wetzel, Silvia Bulgheresi, Stefan Sommer, and Tina Enders. In addition, we would like to thank the crew of the Meteor cruise M192, as well as the Carrie Bow Cay Laboratory, the Heron Island Research Station, the HYDRA Institute Elba, the Mediterranean Institute for Advanced Studies, the Lee Stocking Island Research Station, the Little Darby Island Research Station, the Lizard Island Research Station, and the Okinawa Institute of Science and Technology and their staff for supporting our sampling campaigns.

## Author Contributions

EK and ND conceived the study and designed the workflows with help of KW, ICO, AM, GDA and APF. EK, KW, AM, ICO, APF and GDA analyzed the data. EK and AM interpreted the results. EK drafted the manuscript and EK, AM, ALDO, APF, HS and ND edited the manuscript. All authors provided revisions.

## Competing Interests statement

The authors declare no competing interests.

## Notes

### Competing Interest Statement

The authors have declared no competing interest.

