## Supplementary Figures for "Chemosynthetic Symbioses as Hidden Hubs of DMSP and Organosulfur Cycling in Marine Sediments"

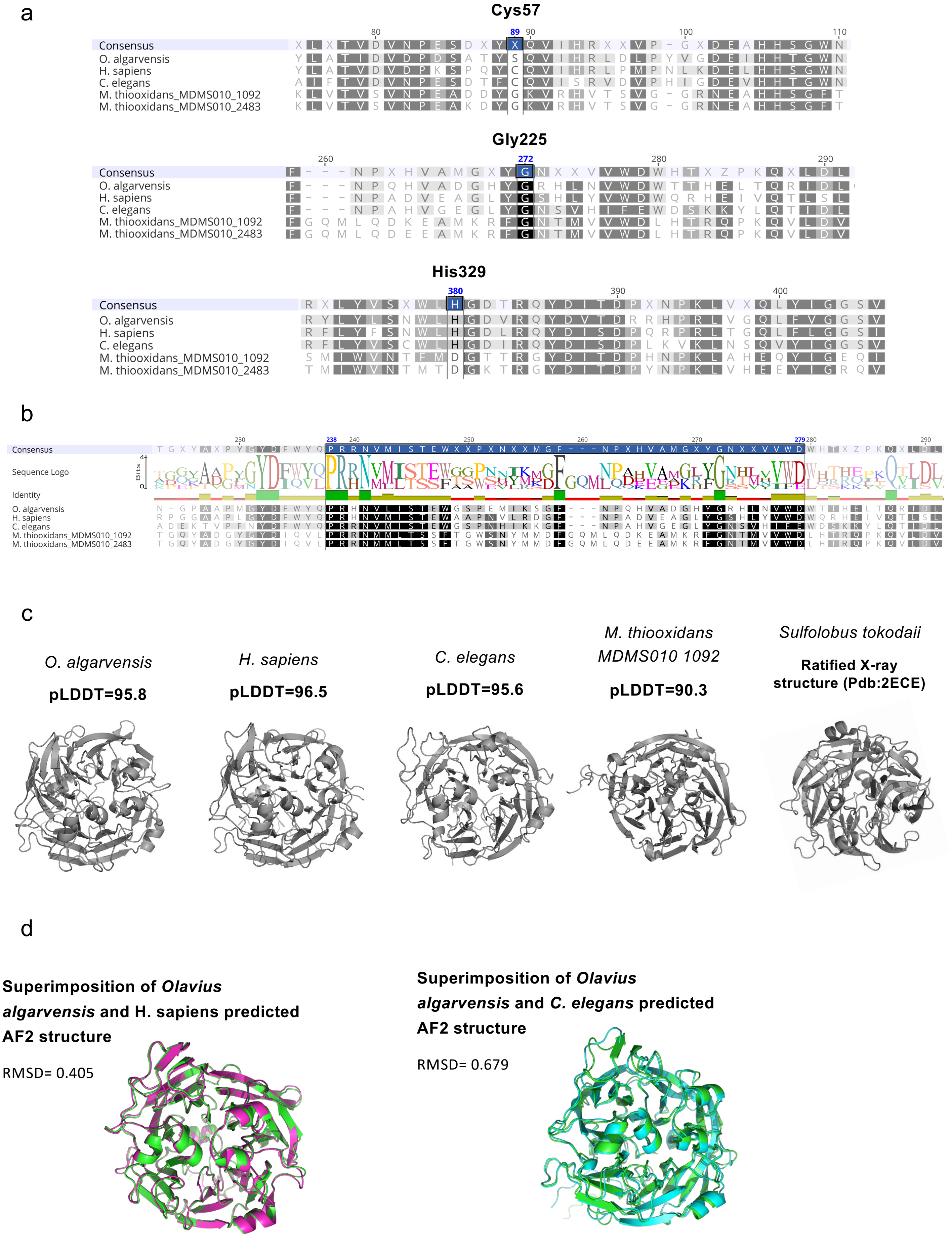


**Supplementary Fig. S1. Conservation and predicted structure of *O. algarvensis* MtoX compared with bacterial and animal homologs.** **(a)** Amino acid alignments of *O. algarvensis* MtoX with *H. sapiens* SELENBP1, *C. elegans* MTOX, and *Methylophaga thiooxidans* MtoX show conservation of two catalytic residues (G272 and H380, blue) known to coordinate the catalytic copper center, while H89 is replaced by serine. **(b)** Sequence logo highlighting conservation of the WD40-like β-propeller motif characteristic of the MtoX/SELENBP1 family. **(c)** AlphaFold2-predicted tertiary structures for all homologs display the canonical eight-bladed β-propeller fold. **(d)** Structural overlays of *O. algarvensis* MtoX with *H. sapiens* SELENBP1 (left) and *M. thiooxidans* MtoX (right) show strong overall structural conservation despite the active-site substitution, suggesting potential retention of methanethiol-oxidase-like function.

a

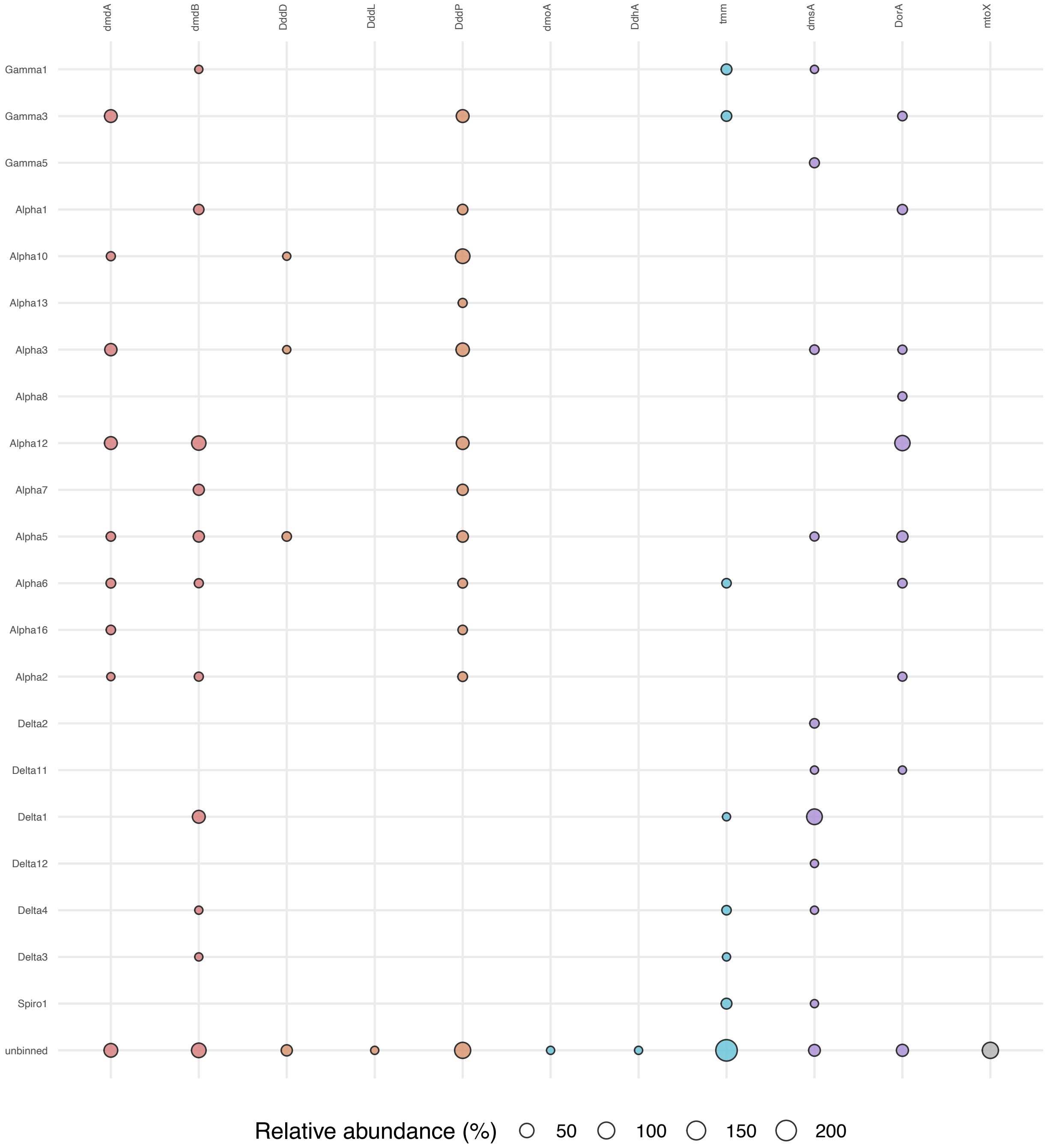


**Figure S2. (a) Distribution of DMSP-, DMS-, and MeSH-cycling genes across symbiont lineages in gutless oligochaetes.** Bubble plot showing the relative abundance of organosulfur metabolism genes detected on contigs assigned to major symbiont lineages (Gamma-, Delta-, Alpha-proteobacteria, Spirochaetes, and unbinned sequences) across 60 gutless oligochaete metagenomes. Bubble size corresponds to the relative abundance (%) of each gene per lineage. **(b) Symbiont-gene enrichment analysis based on Z-scores.** Z-scores were calculated from standardized residuals of a chi-squared test to assess whether specific symbiont lineages are enriched or depleted in particular organosulfur metabolism genes. Positive values (purple) indicate significant enrichment. Gamma3, several Deltaproteobacteria, and multiple Alphaproteobacteria lineages show strong enrichment for distinct gene sets, consistent with their specialized functional roles.

b

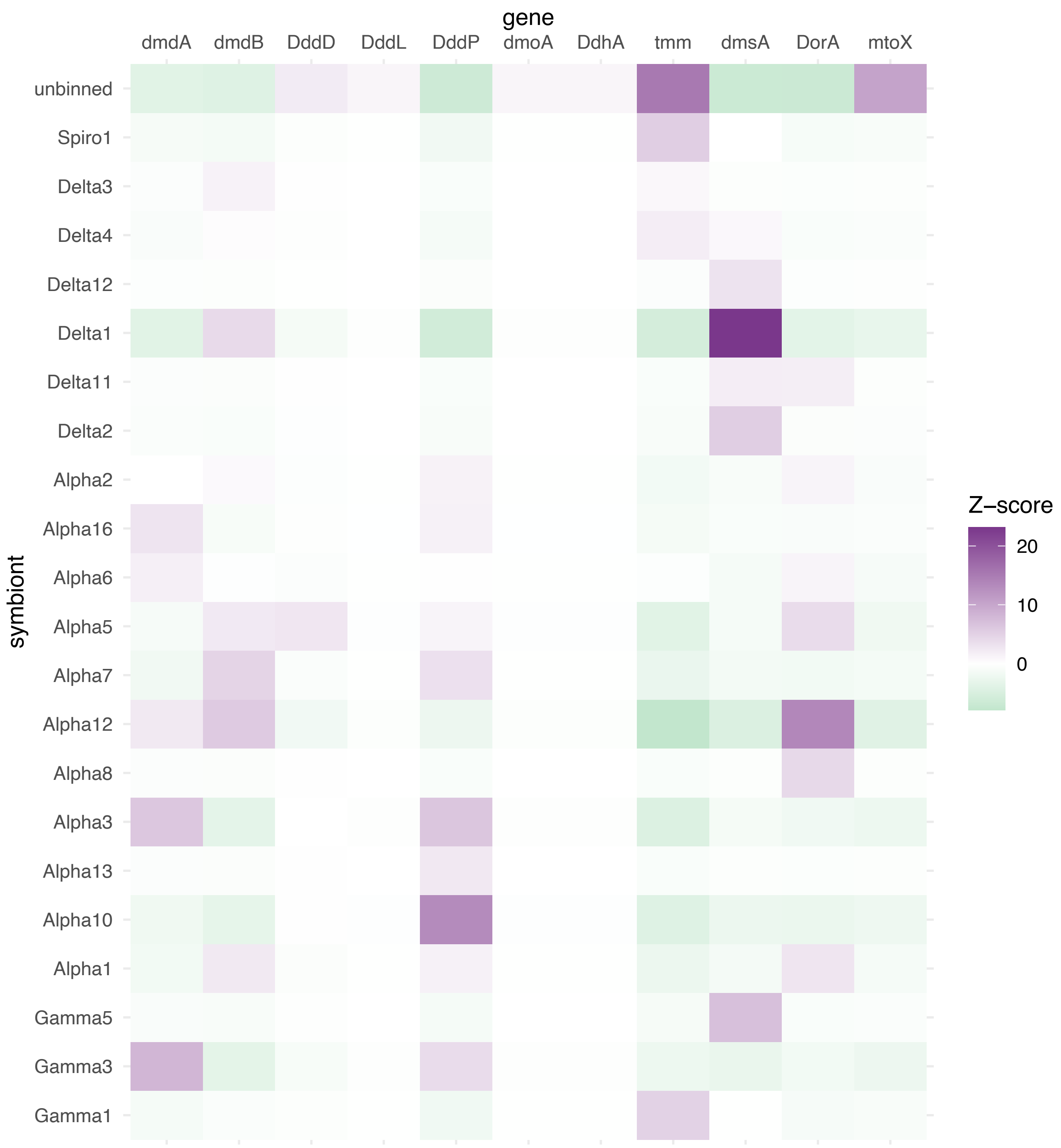
gene
dmdA
dmdB
DddD
DddL
DddP
dmoA
DdhA
tmm
dmsA
DorA
mtoX
symbiont
unbinned
Spiro1
Delta3
Delta4
Delta12
Delta1
Delta11
Delta2
Alpha2
Alpha16
Alpha6
Alpha5
Alpha7
Alpha12
Alpha8
Alpha3
Alpha13
Alpha10
Alpha1
Gamma5
Gamma3
Gamma1
Z−score
20
10
0

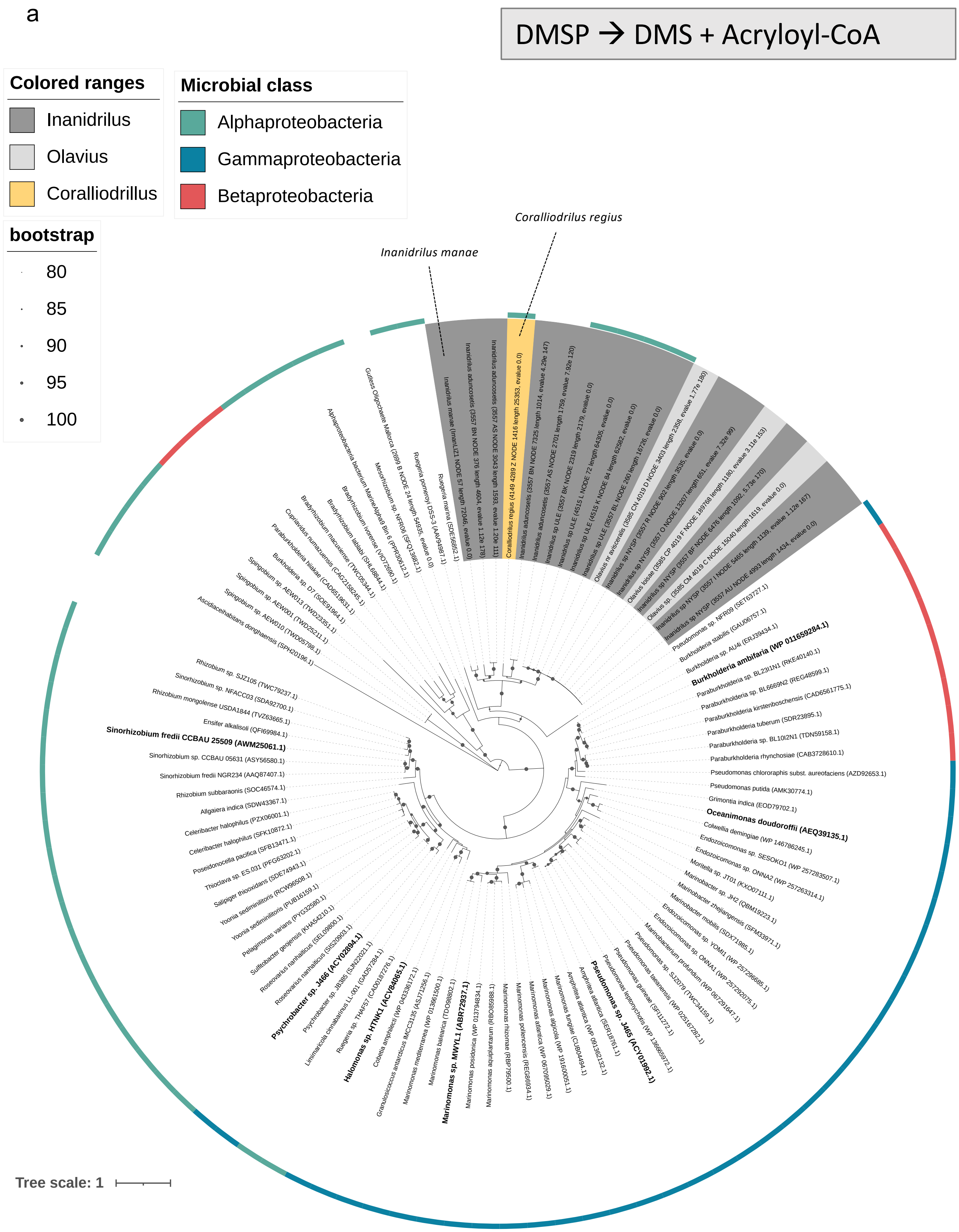


**Figure S3. DMSP lyase *dddD* detected in gutless-oligochaete metagenomes.**
**(a) Maximum-likelihood tree of DddD proteins** including assembly-derived sequences from this study and reference sequences compiled from Chiou et al. 2023 and Liu et al. 2022, supplemented with GenBank/UniProt entries. Tip labels end with the accession number, host species are indicated for assembly-derived sequences, and where available, taxonomic annotations are shown in the outer ring. The tree also shows previously ratified proteins (bold). Light and dark grey shadings indicate metagenomes from gutless oligochaetes (*Olavius*/*Inanidrillus*) and yellow shadings indicate metagenomes from gut-bearing representatives. Protein sequences were aligned with Clustal Omega and the tree inferred with IQ-TREE v1.6.12 using ModelFinder. Branch supports was obtained with the ultrafast bootstrap (Hoang et al., 2018) from 1000 replicates, displayed for nodes with ≥80% support; scale bar shows substitutions per site. Samples that were investigated in the gene neighborhood analysis are indicated with a line and the name of the host. **(b) Gene neighborhoods** for representative *dddD*-positive contigs from gutless-oligochaete metagenomes and reference genomes (≤10 kb total span; scale bar = 1 kb). Co-localized genes are annotated. Arrows denote gene orientation.

b

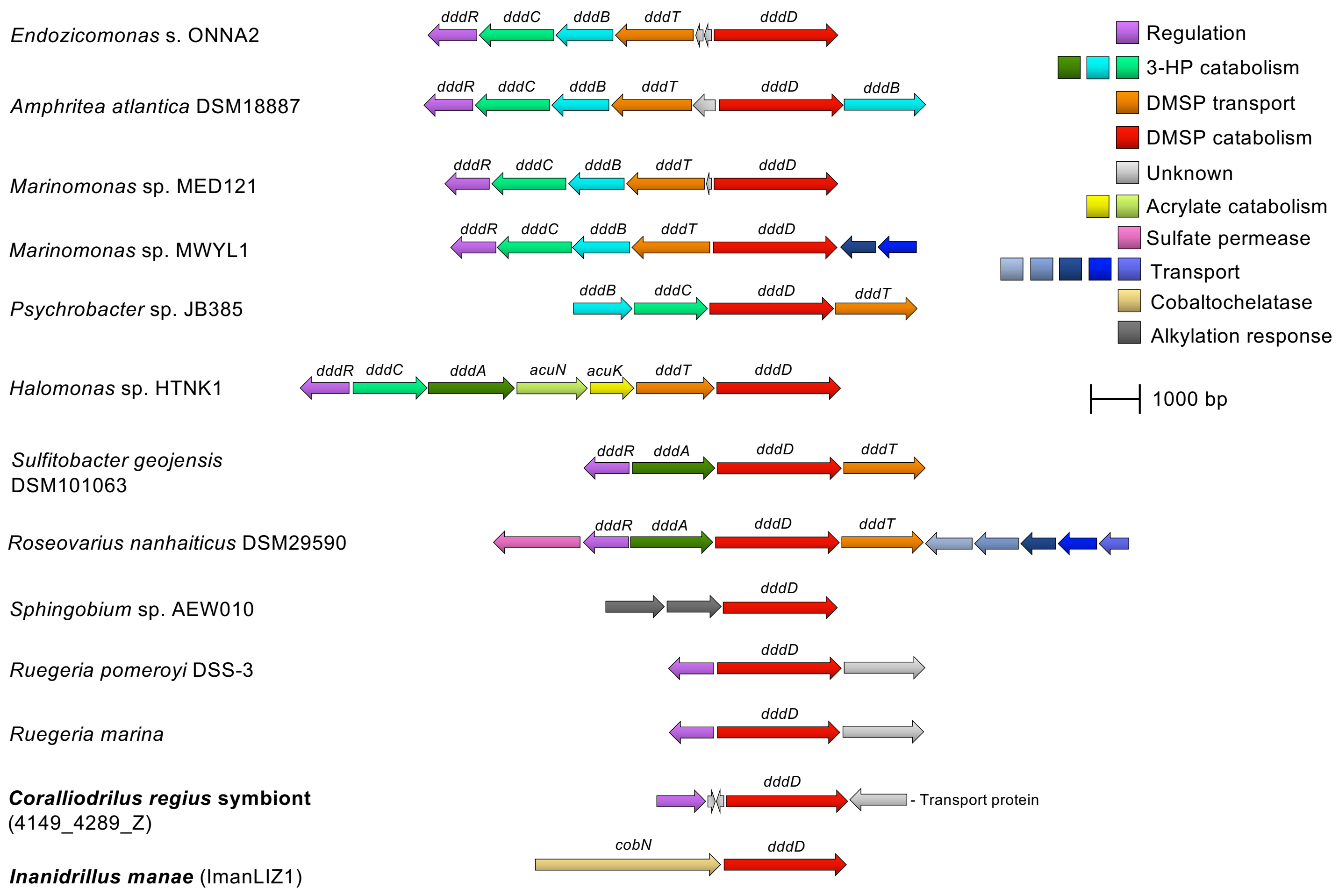
Endozicomonas s. ONNA2
dddR dddC dddB dddT dddD
Amphritea atlantica DSM18887
dddR dddC dddB dddT dddD dddB
Marinomonas sp. MED121
dddR dddC dddB dddT dddD
Marinomonas sp. MWYL1
dddR dddC dddB dddT dddD
Psychrobacter sp. JB385
dddB dddC dddD dddT
Halomonas sp. HTNK1
dddR dddC dddA acuN acuK dddT dddD
Sulfitobacter geojensis DSM101063
dddR dddA dddD dddT
Roseovarius nanhaiticus DSM29590
dddR dddA dddD dddT
Sphingobium sp. AEW010
dddD
Ruegeria pomeroyi DSS-3
dddD
Ruegeria marina
dddD
Coralliodrilus regius symbiont (4149_4289_Z)
dddD
- Transport protein
Inanidrillus manae (ImanLIZ1)
cobN dddD
Regulation
3-HP catabolism
DMSP transport
DMSP catabolism
Unknown
Acrylate catabolism
Sulfate permease
Transport
Cobaltochelatase
Alkylation response
1000 bp

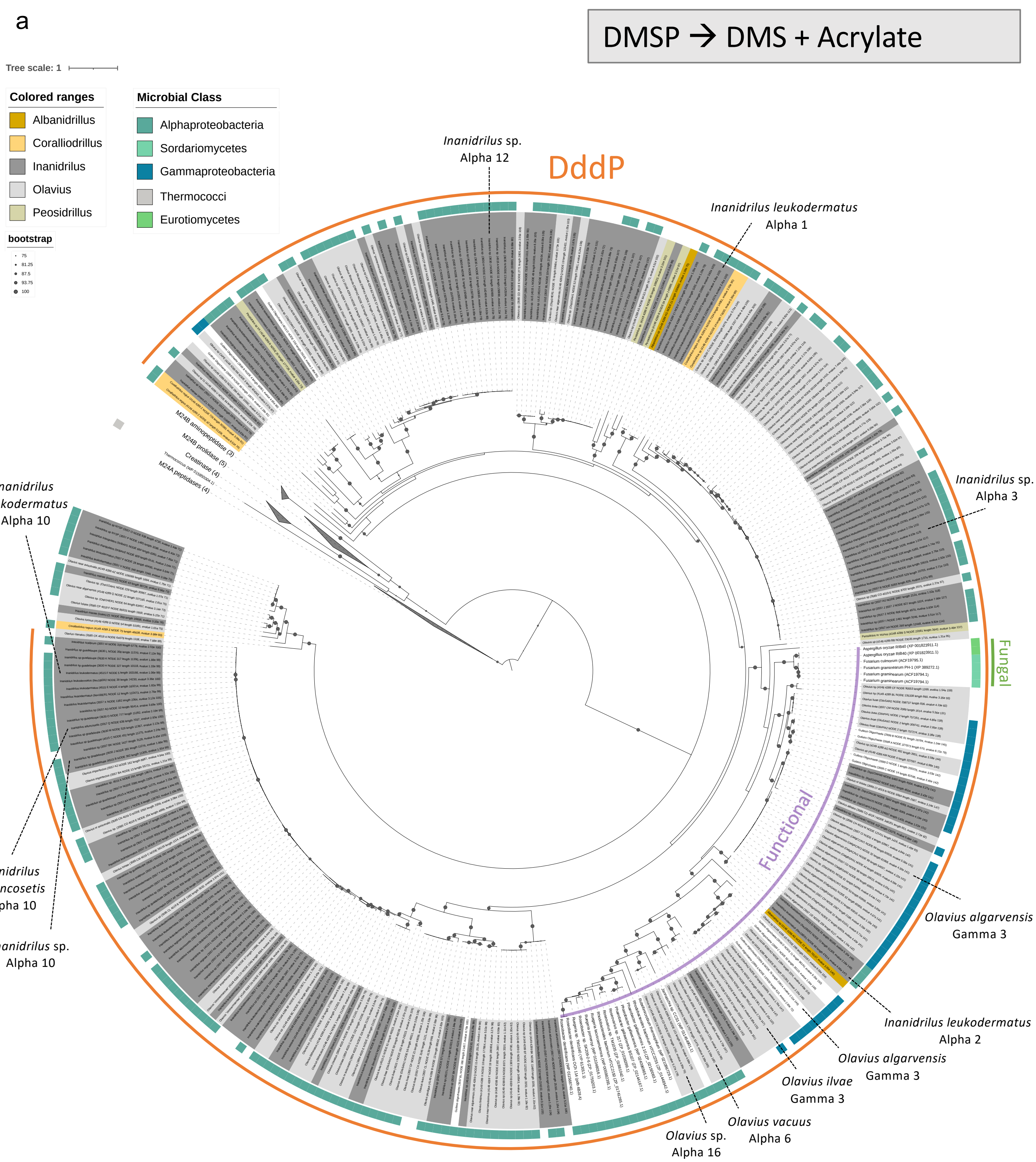


**Figure S4. DMSP lyase *dddP* in gutless-oligochaete metagenomes.**
**(a) Maximum-likelihood phylogeny of DddP.** Assembly-derived DddP sequences from gutless-oligochaete metagenomes are placed among curated references. M24A peptidases (and representative M24B peptidase/prolidase/creatinase proteins) are included as outgroup/context. The clade that contains DddP homologs whose function has been confirmed biochemically is highlighted with a purple line. Fungal DddP homologs are indicated as a distinct clade (marked green). Light and dark grey shadings indicate metagenomes from gutless oligochaetes (*Olavius*/*Inanidrillus*) and yellow, brownish shadings indicate metagenomes from gut-bearing representatives. The outer ring denotes microbial class; tip labels end with accession numbers. Proteins were aligned with Clustal Omega (Geneious Prime v2021.0.3). The tree was inferred with IQ-TREE v1.6.12, ModelFinder model selection, and ultrafast bootstrap (UFBoot) = 1000. Support values are shown for nodes ≥75%; scale bar = substitutions per site. Reference sequences were used from Todd et al. (2009) and Wang et al. Samples that were investigated in the gene neighborhood analysis are indicated with a line and the name of the host/symbiont. (2015). **(b) Gene neighbourhoods.** Representative *dddP*-positive contigs from gutless-oligochaete assemblies and selected references are shown (≤10 kb span; 1 kb scale). Arrows indicate gene orientation. (Neighbourhood gene calls are from this study's contig-level inspection.)

b

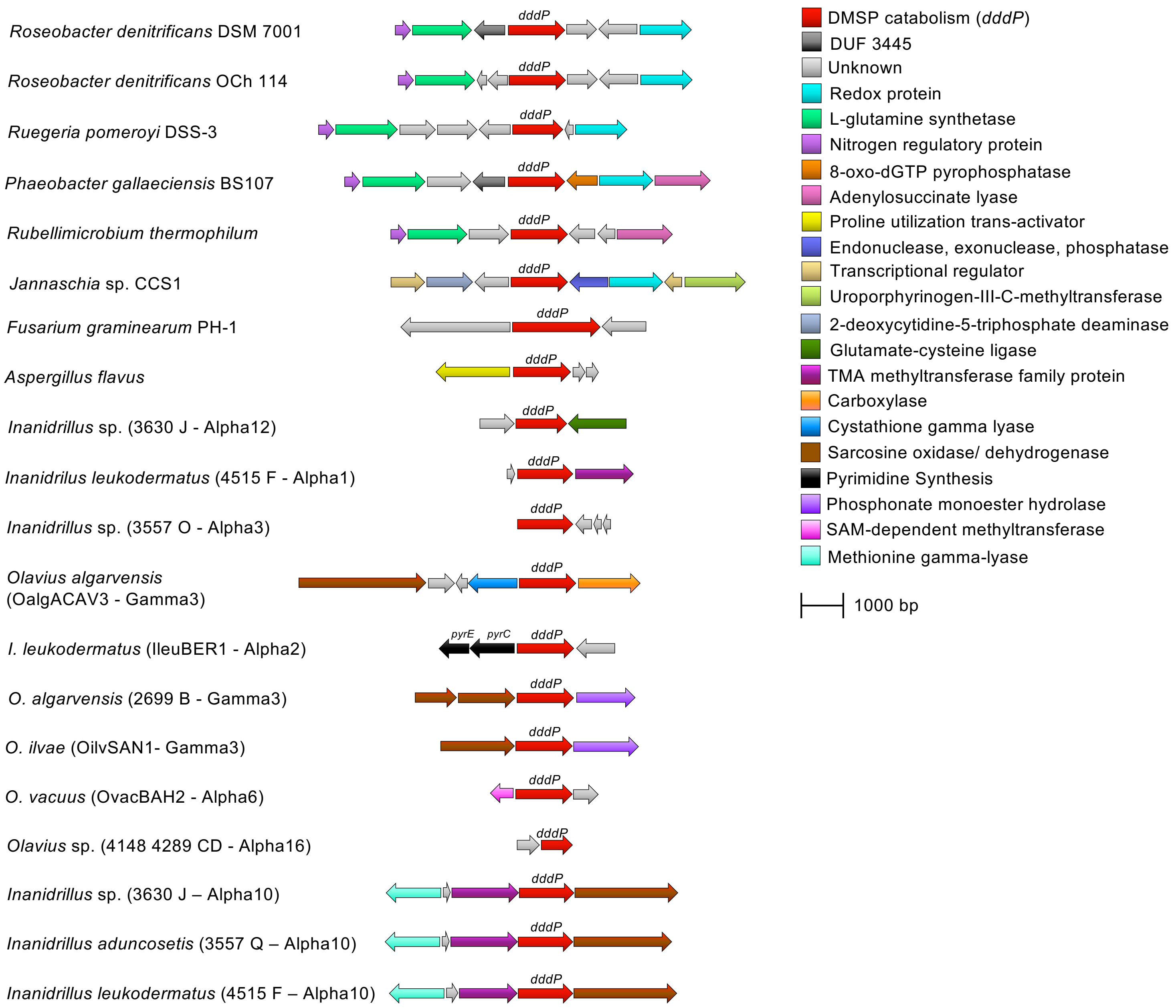

*Roseobacter denitrificans* DSM 7001
*Roseobacter denitrificans* OCh 114
*Ruegeria pomeroyi* DSS-3
*Phaeobacter gallaeciensis* BS107
*Rubellimicrobium thermophilum*
*Jannaschia* sp. CCS1
*Fusarium graminearum* PH-1
*Aspergillus flavus*
*Inanidrillus* sp. (3630 J - Alpha12)
*Inanidrilus leukodermatus* (4515 F - Alpha1)
*Inanidrillus* sp. (3557 O - Alpha3)
*Olavius algarvensis*
(OalgACAV3 - Gamma3)
*I. leukodermatus* (IleuBER1 - Alpha2)
*O. algarvensis* (2699 B - Gamma3)
*O. ilvae* (OilvSAN1- Gamma3)
*O. vacuus* (OvacBAH2 - Alpha6)
*Olavius* sp. (4148 4289 CD - Alpha16)
*Inanidrillus* sp. (3630 J – Alpha10)
*Inanidrillus aduncosetis* (3557 Q – Alpha10)
*Inanidrillus leukodermatus* (4515 F – Alpha10)
*dddP*
*pyrE*
*pyrC*
DMSP catabolism (*dddP*)
DUF 3445
Unknown
Redox protein
L-glutamine synthetase
Nitrogen regulatory protein
8-oxo-dGTP pyrophosphatase
Adenylosuccinate lyase
Proline utilization trans-activator
Endonuclease, exonuclease, phosphatase
Transcriptional regulator
Uroporphyrinogen-III-C-methyltransferase
2-deoxycytidine-5-triphosphate deaminase
Glutamate-cysteine ligase
TMA methyltransferase family protein
Carboxylase
Cystathione gamma lyase
Sarcosine oxidase/ dehydrogenase
Pyrimidine Synthesis
Phosphonate monoester hydrolase
SAM-dependent methyltransferase
Methionine gamma-lyase
1000 bp

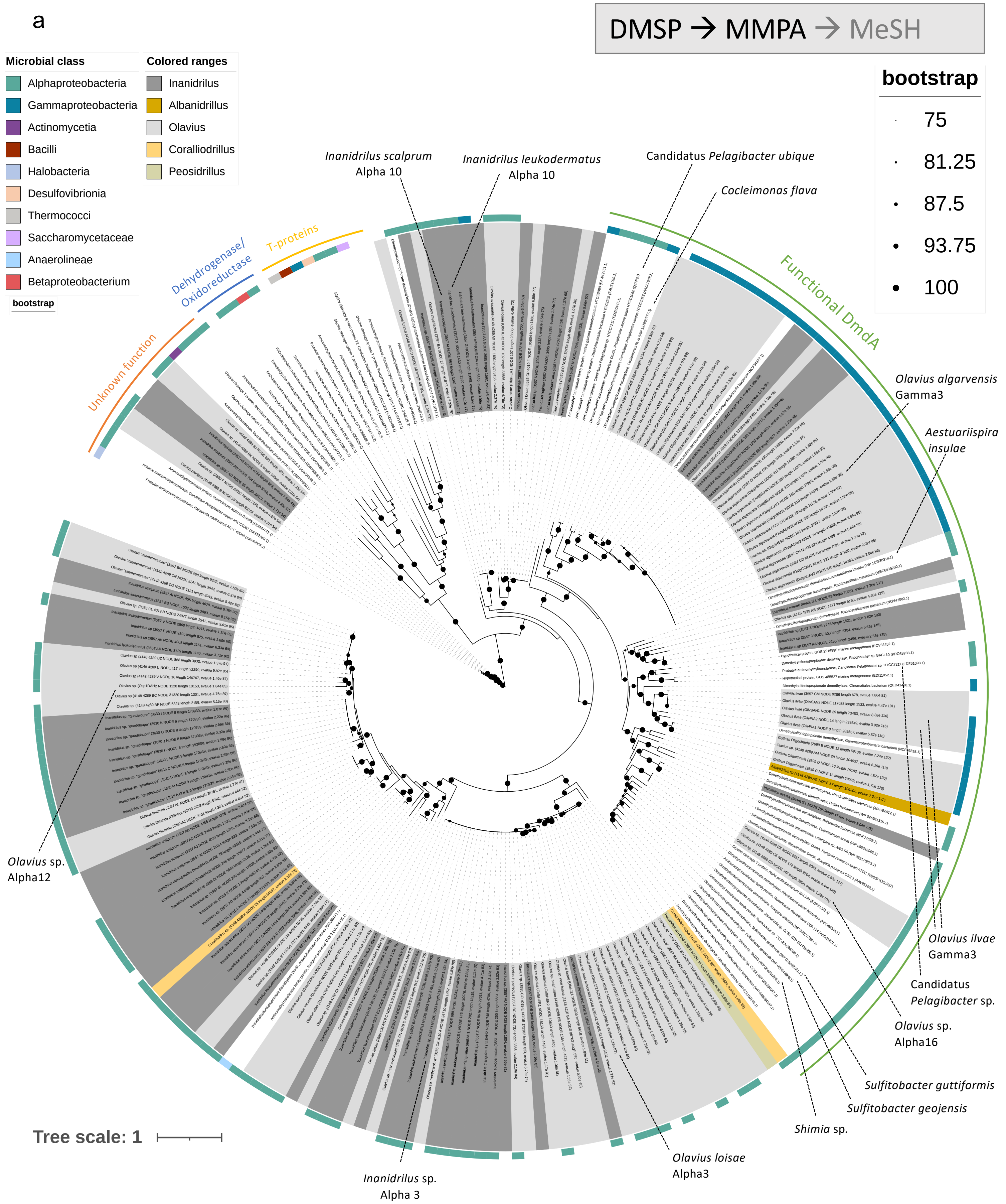


**Figure S5. DMSP demethylase *dmdA* in gutless-oligochaete metagenomes**

**(a) Maximum-likelihood phylogeny of DmdA.** Assembly-derived DmdA sequences from gutless-oligochaete metagenomes are placed among curated references. The reference set was compiled from Reisch et al. 2008 and Hernández et al. 2020. Following Reisch et al., proteins of unknown function (non-DmdA homologs) were included as outgroup, together with GcvT (glycine-cleavage T-proteins) and selected dehydrogenases/oxidoreductases to anchor the boundary between *bona fide* DmdA and related enzymes. Tip labels end with accession numbers; the outer ring indicates microbial class. Proteins were aligned with Clustal Omega (Geneious Prime v2021.0.3); the tree was inferred with IQ-TREE v1.6.12 using ModelFinder and UFBoot=1000; support values are shown for nodes ≥75%; scale bar = substitutions/site. Sequences that fall within the "unknown-function/non-DmdA" clade are flagged and not scored as *dmdA* in prevalence summaries. Samples that were investigated in the gene neighborhood analysis are indicated with a line and the name of the microorganism or host/symbiont. **(b) Gene neighbourhoods.** Representative *dmdA*-positive contigs from gutless-oligochaete assemblies and selected references (≤10 kb span; 1 kb scale) are shown. In *Pelagibacter* and two alphaproteobacterial symbiont-derived contigs, *dmdA* co-localizes with *dmdB* and often hydratase/hydrolase-like and *dmdC* genes, consistent with a contiguous *dmdABC* module. In other contigs, *dmdA* lies near a candidate acrylyl-CoA reductase (AcuI/Acr-like). One *Olavius algarvensis* neighbourhood (OalgBSAN1 - Gamma3) also shows proximity to an enoyl-CoA

b

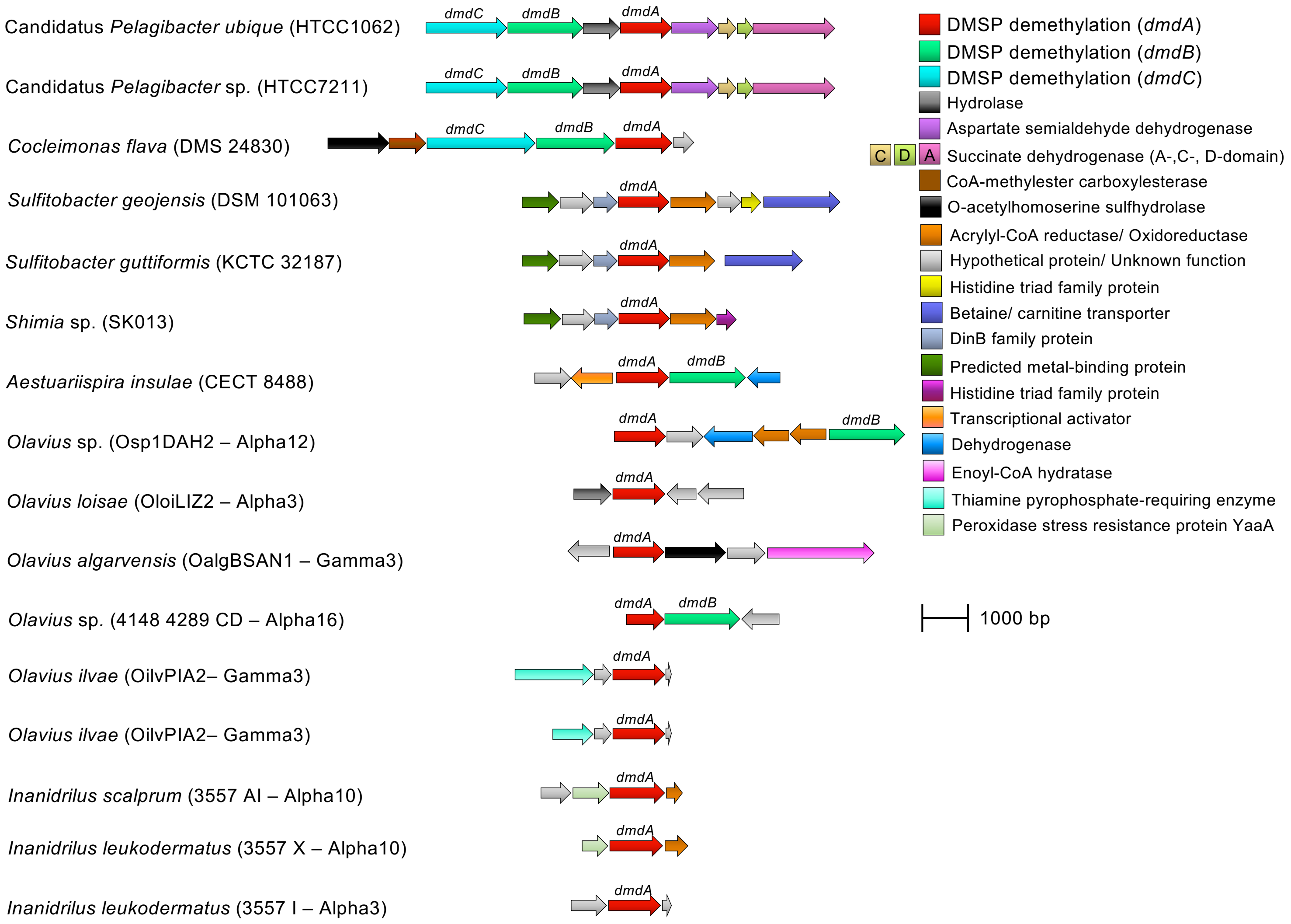
Candidatus *Pelagibacter ubique* (HTCC1062)
dmdC
dmdB
dmdA
Candidatus *Pelagibacter* sp. (HTCC7211)
dmdC
dmdB
dmdA
*Cocleimonas flava* (DMS 24830)
dmdC
dmdB
dmdA
*Sulfitobacter geojensis* (DSM 101063)
dmdA
*Sulfitobacter guttiformis* (KCTC 32187)
dmdA
*Shimia* sp. (SK013)
dmdA
*Aestuariispira insulae* (CECT 8488)
dmdA
dmdB
*Olavius* sp. (Osp1DAH2 – Alpha12)
dmdA
dmdB
*Olavius loisae* (OloiLIZ2 – Alpha3)
dmdA
*Olavius algarvensis* (OalgBSAN1 – Gamma3)
dmdA
*Olavius* sp. (4148 4289 CD – Alpha16)
dmdA
dmdB
*Olavius ilvae* (OilvPIA2– Gamma3)
dmdA
*Olavius ilvae* (OilvPIA2– Gamma3)
dmdA
*Inanidrilus scalprum* (3557 AI – Alpha10)
dmdA
*Inanidrilus leukodermatus* (3557 X – Alpha10)
dmdA
*Inanidrilus leukodermatus* (3557 I – Alpha3)
dmdA
DMSP demethylation (*dmdA*)
DMSP demethylation (*dmdB*)
DMSP demethylation (*dmdC*)
Hydrolase
Aspartate semialdehyde dehydrogenase
C D A Succinate dehydrogenase (A-,C-, D-domain)
CoA-methylester carboxylesterase
O-acetylhomoserine sulfhydrolase
Acrylyl-CoA reductase/ Oxidoreductase
Hypothetical protein/ Unknown function
Histidine triad family protein
Betaine/ carnitine transporter
DinB family protein
Predicted metal-binding protein
Histidine triad family protein
Transcriptional activator
Dehydrogenase
Enoyl-CoA hydratase
Thiamine pyrophosphate-requiring enzyme
Peroxidase stress resistance protein YaaA
1000 bp

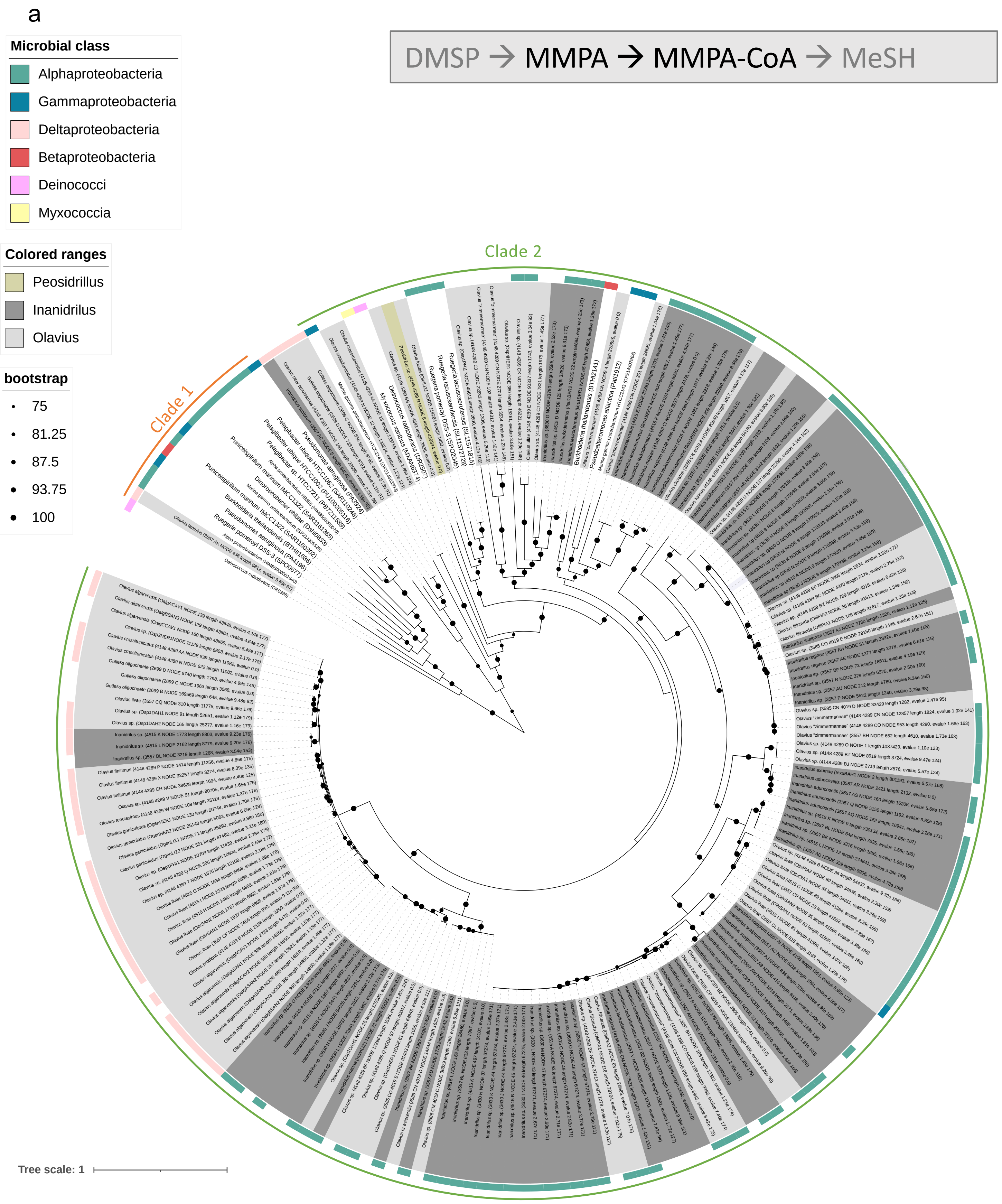


**Figure S6. MMPA–CoA ligase *dmdB* in gutless-oligochaete metagenomes**
**Maximum-likelihood phylogeny of DmdB.** Assembly-derived DmdB sequences from gutless-oligochaete metagenomes are shown alongside a curated reference set emphasizing the two canonical DmdB clades (clade1/clade2) reported previously (Bullock et al. 2014). Tip labels end with accession numbers, and the outer ring indicates microbial class. Proteins were aligned with Clustal Omega (Geneious Prime v2021.0.3); the tree was inferred with IQ-TREE v1.6.12 using ModelFinder and ultrafast bootstrap (UFBoot) = 1000; support values are shown for nodes ≥60%; scale bar = substitutions/site. Reference sequences were drawn from Bullock et al., 2014.

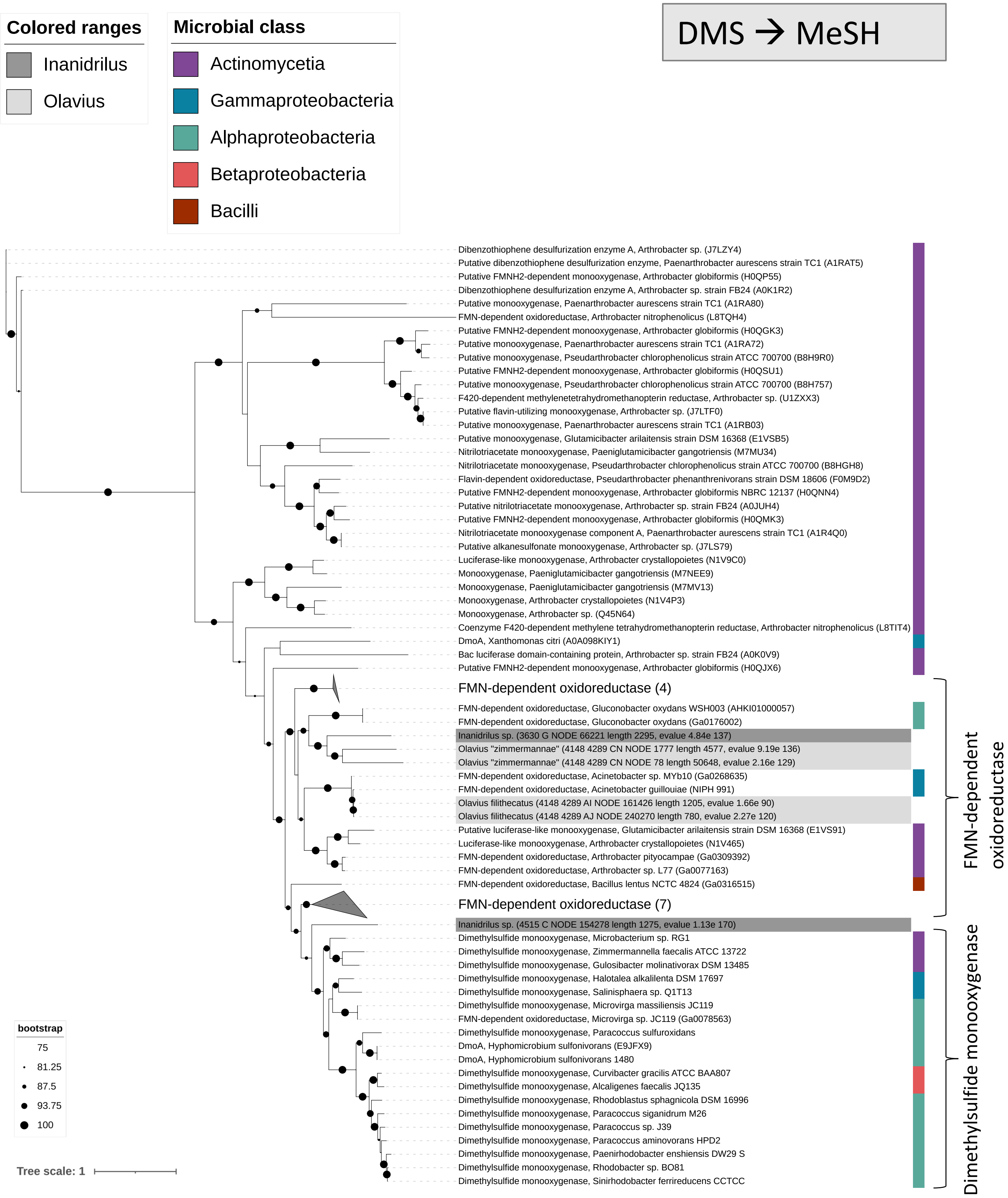


**Figure S7. DMS monooxygenase A-subunit *dmoA* in gutless-oligochaete metagenomes.**
Maximum-likelihood phylogeny of DmoA and related $FMNH_2$-dependent oxidoreductases. Assembly-derived candidates from gutless-oligochaete metagenomes were placed among curated references (including sequences from Hammers et al. 2020). Dibenzothiophene desulfurization enzyme was used as an outgroup. Tip labels end with accession numbers. Proteins were aligned with Clustal Omega (Geneious Prime v2021.0.3); the tree was inferred with IQ-TREE v1.6.12 using ModelFinder; UFBoot = 1000; support values are shown for nodes ≥75%; scale bar indicates substitutions/site.

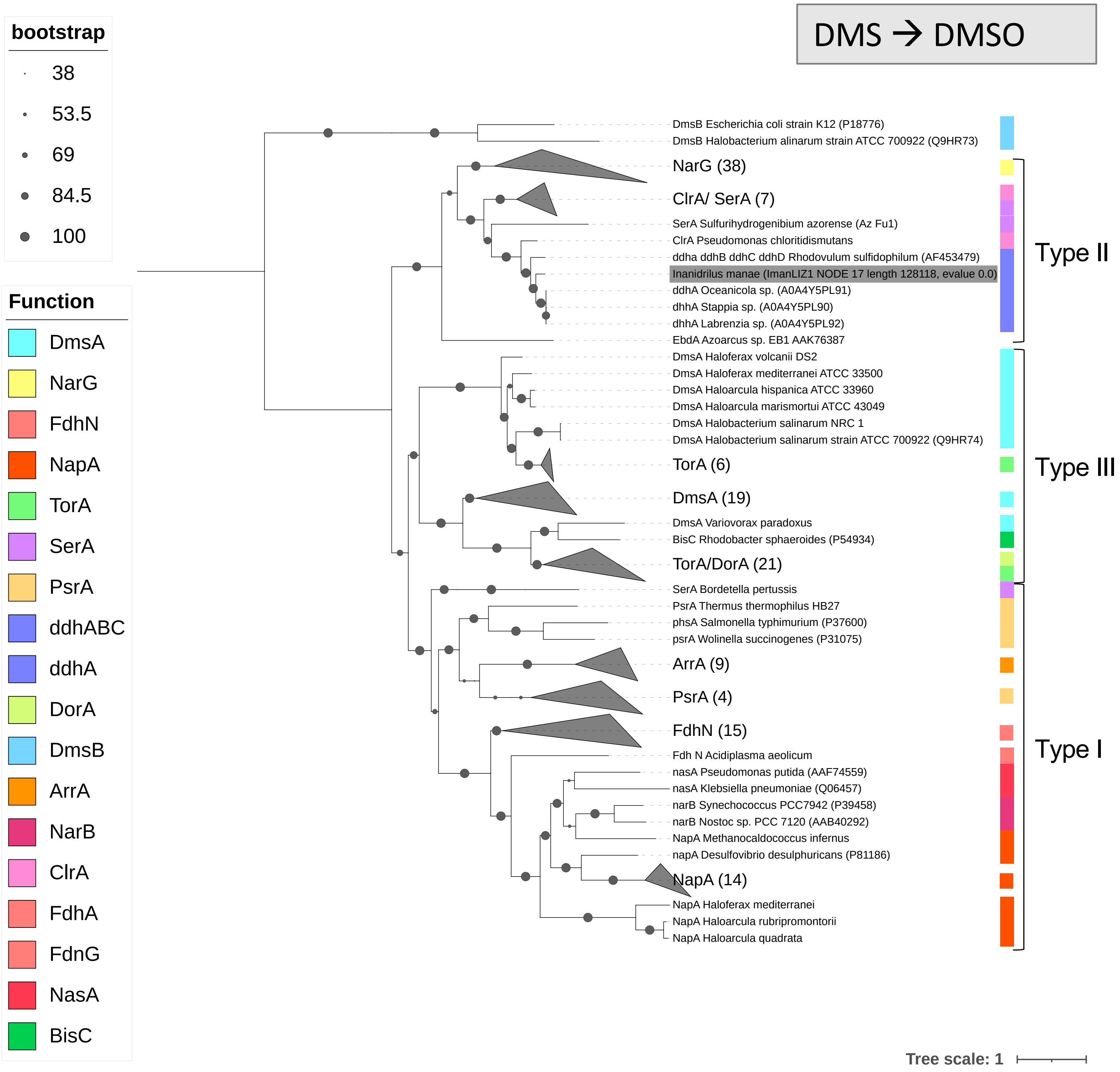


**Figure S8. *ddhA* (DMS dehydrogenase, catalytic subunit) in gutless-oligochaete metagenomes. Maximum-likelihood phylogeny of DdhA within the DMSO-reductase family.** Assembly-derived DdhA candidates from gutless-oligochaete metagenomes are placed among curated references from *Rhodovulum sulfidophilum* and other DMSO-reductase family members (NarG/NapA/DmsA/SerA/EbdA etc.) from Yamazaki et al. 2020 and McDevitt et al.2002. The iron-sulfur subunit DmsB was included as an outgroup; additional DMSO-reductase family sequences were added as closely related context. Proteins were aligned with Clustal Omega (Geneious Prime v2021.0.3); tree was inferred with IQ-TREE v1.6.12 using ModelFinder; node support is UFBoot from 1000 replicates, shown for nodes ≥38%; scale bar = substitutions/site. Tip labels end with accession numbers; DMSO-reductase family members are shown as a bar on the right side. Abbreviations: DmsA/DmsB, periplasmic bacterial DMSO reductase; NarG, respiratory nitrate reductase; FdhA/FdhG/FdhN, formate dehydrogenase; NapA, periplasmic nitrate reductase; TorA, trimethylamine N-oxide (TMAO) reductase; SerA, selenate reductase; PsrA, Polysulfide reductase;DdhA/DdhABC, DMS dehydrogenase; DorA, DMSO reductase; ArrA, respiratory arsenate reductases; NarB/NasA, assimilatory nitrate reductase; ClrA, chlorate reductase; BisC, biotin sulfoxide reductase.

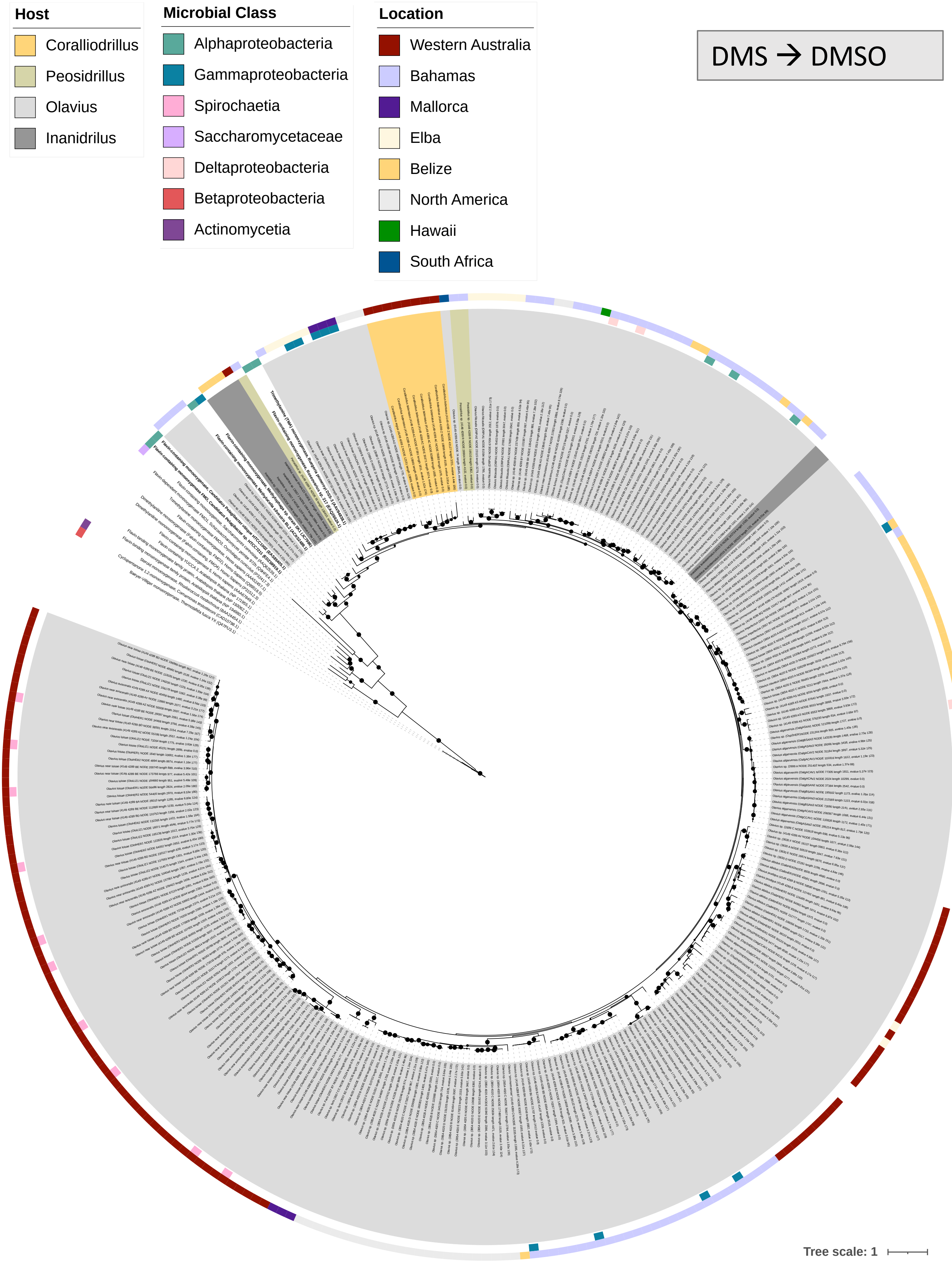


**Figure S9. Maximum-likelihood phylogeny of Tmm and related FMOs.**
Assembly-derived Tmm candidates from gutless-oligochaete metagenomes were placed with curated references (including Chen et al. 2011) and Baeyer-Villiger monooxygenases (BVMO) as outgroup. Light and dark grey shadings indicate metagenomes from gutless oligochaetes (*Olavius*/*Inanidrillus*) and yellow, brownish shadings indicate metagenomes from gut-bearing representatives. Tip labels end with accession numbers. The inner one of the outer ring shows microbial class. The second outer ring shows sampling location of gutless oligochaete metagenomes. Alignment was done with Clustal Omega (Geneious Prime v2021.0.3); tree was inferred with IQ-TREE v1.6.12 with ModelFinder, UFBoot = 1000; support is shown for nodes ≥75%; scale bar = substitutions/site.

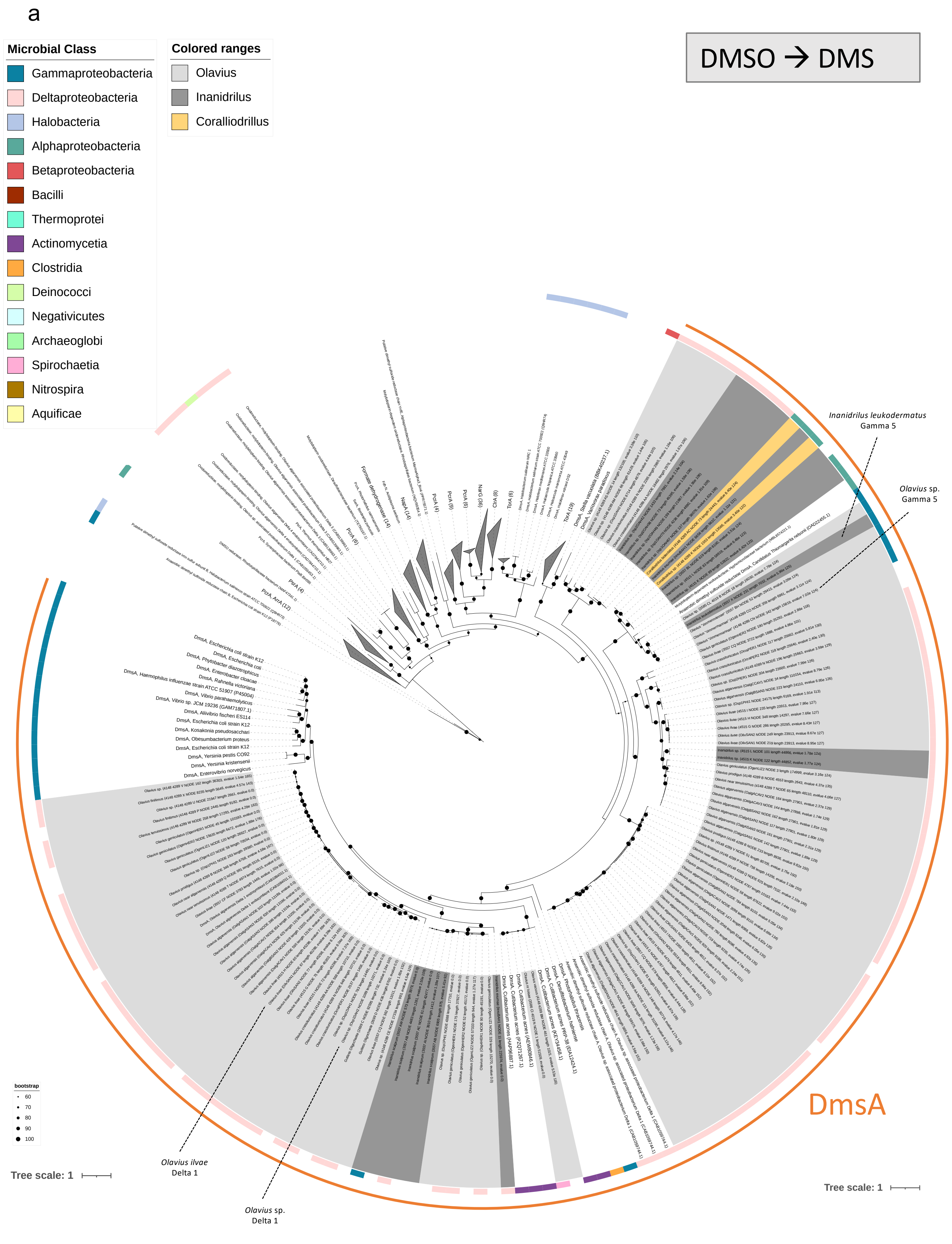


**Figure S10. (a) Maximum-likelihood phylogeny of DmsA in gutless-oligochaete metagenomes within the DMSO-reductase family.** Assembly-derived DmsA sequences from gutless-oligochaete metagenomes were placed among curated references representing the DMSO-reductase superfamily (e.g., NarG/NapA/FdhA/TorA/DorA) from Yamazaki et al. 2020 and McDevitt et al.2002 used as close context. Light and dark grey shadings indicate metagenomes from gutless oligochaetes (*Olavius*/*Inanidrillus*) and yellow shadings from gut-bearing representatives. Proteins were aligned with Clustal Omega (Geneious Prime v2021.0.3); tree was inferred with IQ-TREE v1.6.12 using ModelFinder; node support is UFBoot from 1000 replicates and shown for nodes ≥60%; scale bar = substitutions per site. Tip labels end with accession numbers, and the outer ring indicates microbial class. Samples that were investigated in the gene neighborhood analysis are indicated with a line and the name of the host/symbiont. **(b) Gene neighbourhoods**. Representative *dmsA*-positive contigs from gutless-oligochaete assemblies and selected references (≤10 kb span; 1 kb scale) are shown. *dmsA* co-localized with *dmsB* (Fe-S subunit) and *dmsC* (membrane anchor). Windows ≤10 kb total span; scale bar = 1 kb. Arrows denote gene orientation.

b

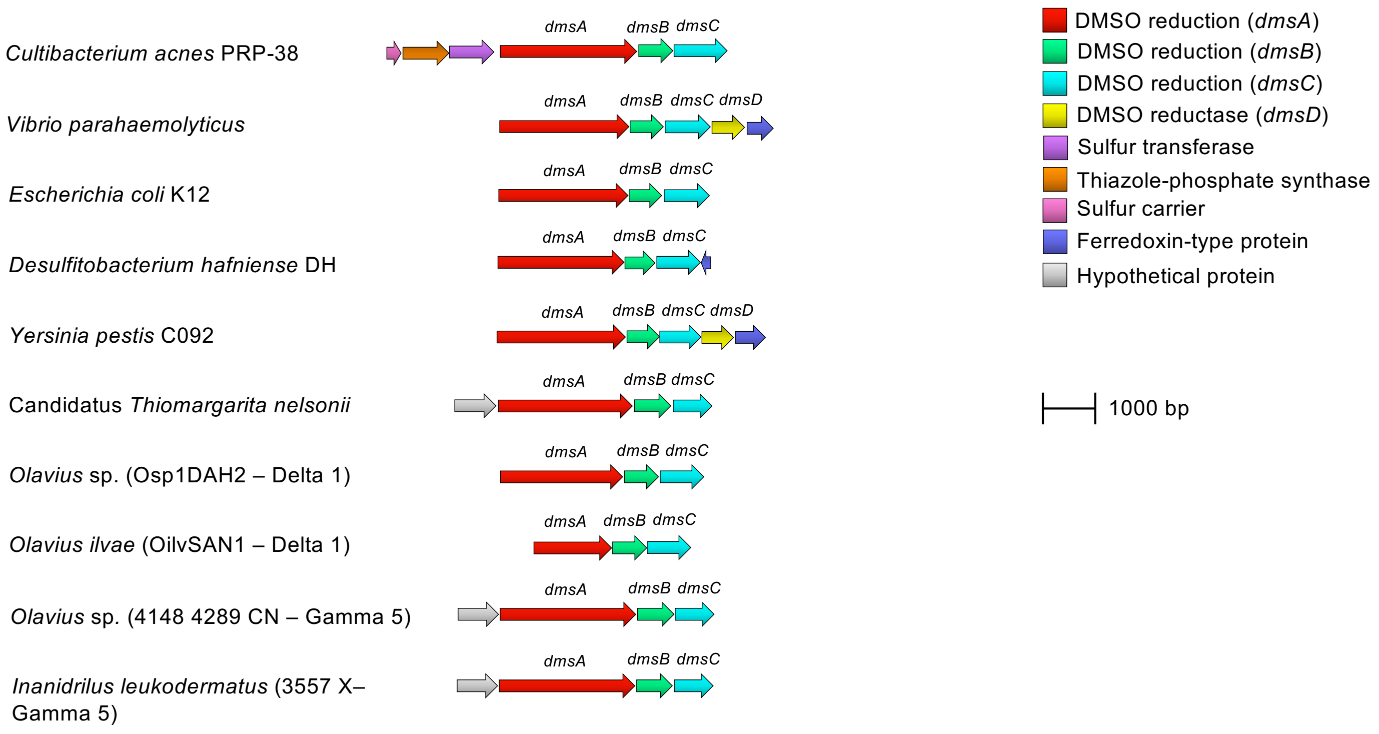
Cultibacterium acnes PRP-38
Vibrio parahaemolyticus
Escherichia coli K12
Desulfitobacterium hafniense DH
Yersinia pestis C092
Candidatus Thiomargarita nelsonii
Olavius sp. (Osp1DAH2 – Delta 1)
Olavius ilvae (OilvSAN1 – Delta 1)
Olavius sp. (4148 4289 CN – Gamma 5)
Inanidrilus leukodermatus (3557 X–Gamma 5)
dmsA
dmsB
dmsC
dmsD
DMSO reduction (dmsA)
DMSO reduction (dmsB)
DMSO reduction (dmsC)
DMSO reductase (dmsD)
Sulfur transferase
Thiazole-phosphate synthase
Sulfur carrier
Ferredoxin-type protein
Hypothetical protein
1000 bp

a

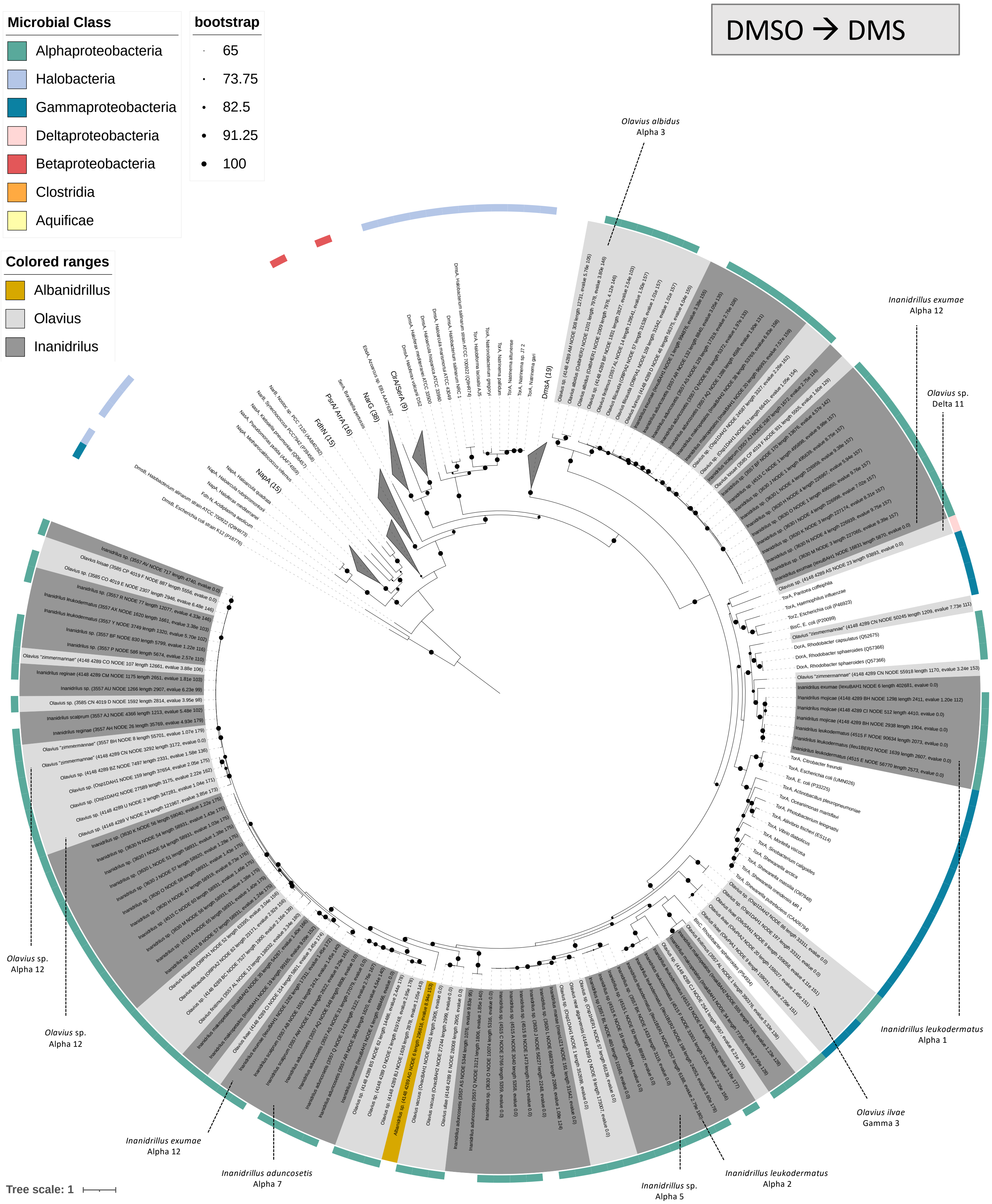


**Figure S11. Periplasmic N-oxide/DMSO reductases (*torA/dorA*) in gutless-oligochaete metagenomes. (a) Maximum-likelihood phylogeny of TorA/DorA within the DMSO-reductase superfamily.** Assembly-derived TorA/DorA candidates from gutless-oligochaete metagenomes were placed among close relatives from the DMSO-reductase family (NarG/NapA/FdhA/DmsA, etc.) for context. DmsB (Fe–S subunit of DmsABC) was used as the outgroup. Proteins were aligned with Clustal Omega (Geneious Prime v2021.0.3); the tree was inferred with IQ-TREE v1.6.12 using ModelFinder; UFBoot = 1000; support values are shown for nodes ≥65%; scale bar = substitutions/site. Tip labels end with accession numbers; an outer ring indicates microbial class. Samples that were investigated in the gene neighborhood analysis are indicated with a line and the name of the host/symbiont. **(b) Gene neighbourhoods**. Representative *torA/dorA*-positive contigs and reference genomes showing *torA/dorA* flanked by canonical partner/regulatory genes (windows ≤10 kb; scale bar = 1 kb). Common cluster features: *torC/dorC* (c-type cytochrome), *torD/dorD* (maturation chaperone), *torS/torR* (regulators), *torT* (periplasmic TMAO sensor), occasional *torY/torZ*, and adjacent Molybdenum cofactor riboswitch. Arrows denote gene orientation.

b

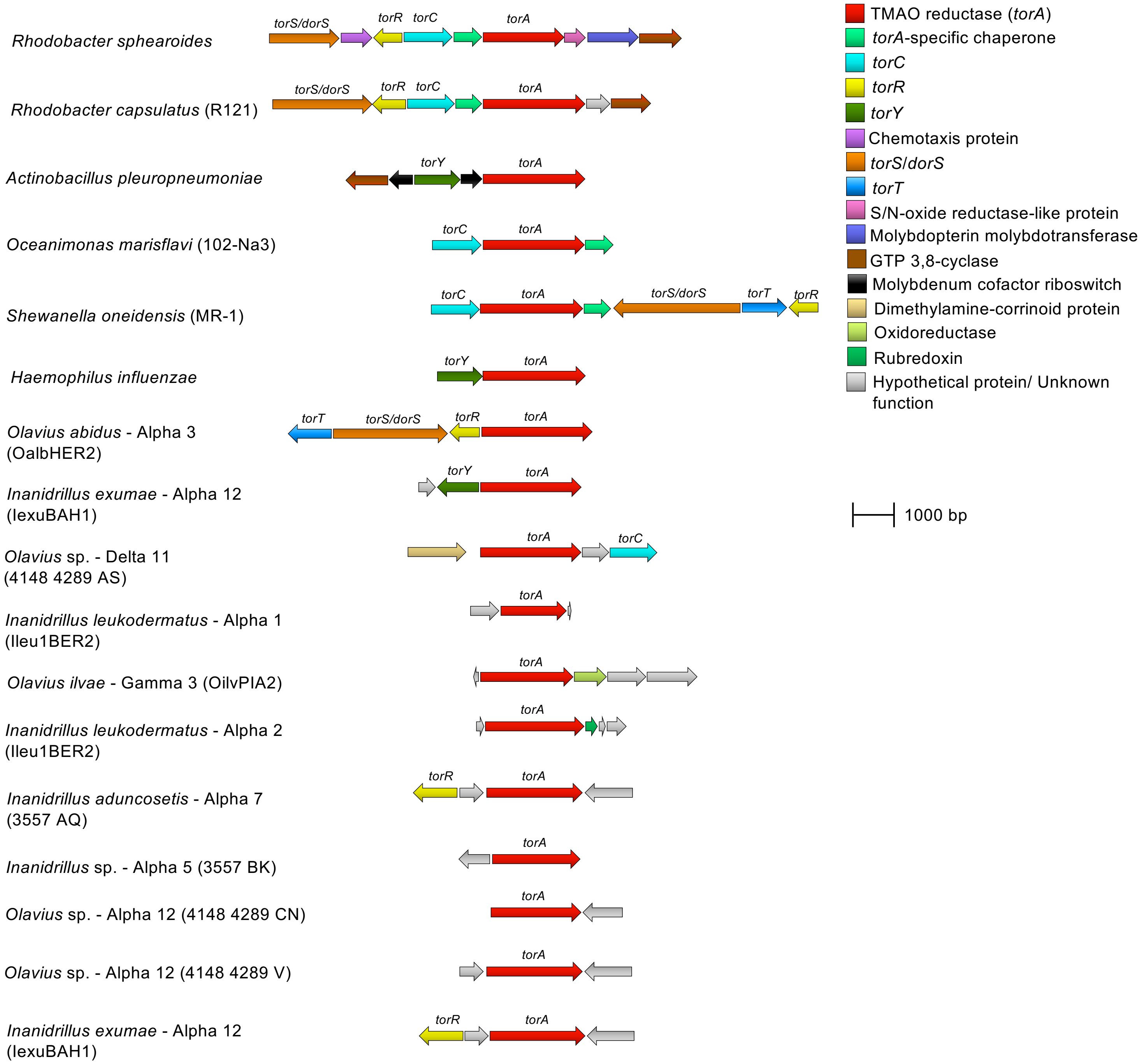
Rhodobacter sphearoides
torS/dorS
torR
torC
torA
Rhodobacter capsulatus (R121)
torS/dorS
torR
torC
torA
Actinobacillus pleuropneumoniae
torY
torA
Oceanimonas marisflavi (102-Na3)
torC
torA
Shewanella oneidensis (MR-1)
torC
torA
torS/dorS
torT
torR
Haemophilus influenzae
torY
torA
Olavius abidus - Alpha 3 (OalbHER2)
torT
torS/dorS
torR
torA
Inanidrillus exumae - Alpha 12 (IexuBAH1)
torY
torA
Olavius sp. - Delta 11 (4148 4289 AS)
torA
torC
Inanidrillus leukodermatus - Alpha 1 (Ileu1BER2)
torA
Olavius ilvae - Gamma 3 (OilvPIA2)
torA
Inanidrillus leukodermatus - Alpha 2 (Ileu1BER2)
torA
Inanidrillus aduncosetis - Alpha 7 (3557 AQ)
torR
torA
Inanidrillus sp. - Alpha 5 (3557 BK)
torA
Olavius sp. - Alpha 12 (4148 4289 CN)
torA
Olavius sp. - Alpha 12 (4148 4289 V)
torA
Inanidrillus exumae - Alpha 12 (IexuBAH1)
torR
torA
TMAO reductase (torA)
torA-specific chaperone
torC
torR
torY
Chemotaxis protein
torS/dorS
torT
S/N-oxide reductase-like protein
Molybdopterin molybdotransferase
GTP 3,8-cyclase
Molybdenum cofactor riboswitch
Dimethylamine-corrinoid protein
Oxidoreductase
Rubredoxin
Hypothetical protein/ Unknown function
1000 bp

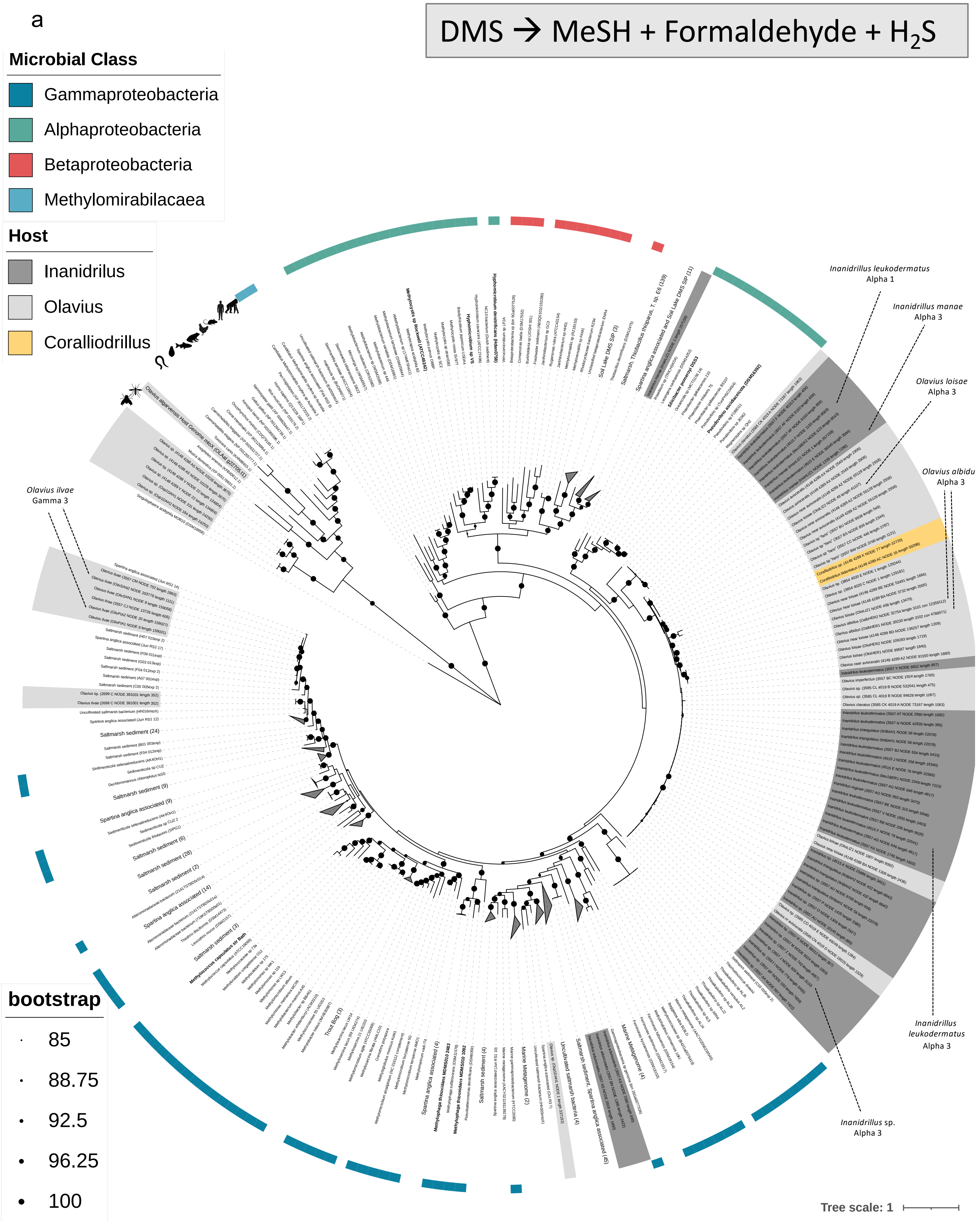


**Figure S12. *mtoX* (methanethiol oxidase; SBP56-family) in gutless-oligochaete metagenomes. (a) Maximum-likelihood phylogeny of MtoX.** Assembly-derived MtoX candidates from gutless-oligochaete metagenomes were placed alongside curated references, including sequences from Eyice et al. (2018) and Kröber et al. (2022). The tree also includes animal methanethiol oxidase homologs (SELENBP1) from e.g. *O. algarvensis*, *C. elegans*, and human, which form a distinct clade from bacterial *mtoX*, confirming their independent evolutionary origin. Tip labels end with accession numbers; the outer ring indicates microbial class. Proteins were aligned with Clustal Omega (Geneious Prime v2021.0.3); tree was inferred with IQ-TREE v1.6.12 using ModelFinder; UFBoot = 1000; support values are shown for nodes ≥75%; scale bar = substitutions/site. Samples that were investigated in the gene neighborhood analysis are indicated with a line and the name of the host/symbiont. **(b) Genomic context (schematic).** Representative reference genomes illustrate the common mtoX-SCO1/senC-mauG arrangement (sometimes SCO1-mauG fusion); gutless-derived *mtoX*-positive contigs are variable/fragmentary and shown where sufficient flanking sequence exists. (Window ≤10 kb; scale bar = 1 kb.)

b

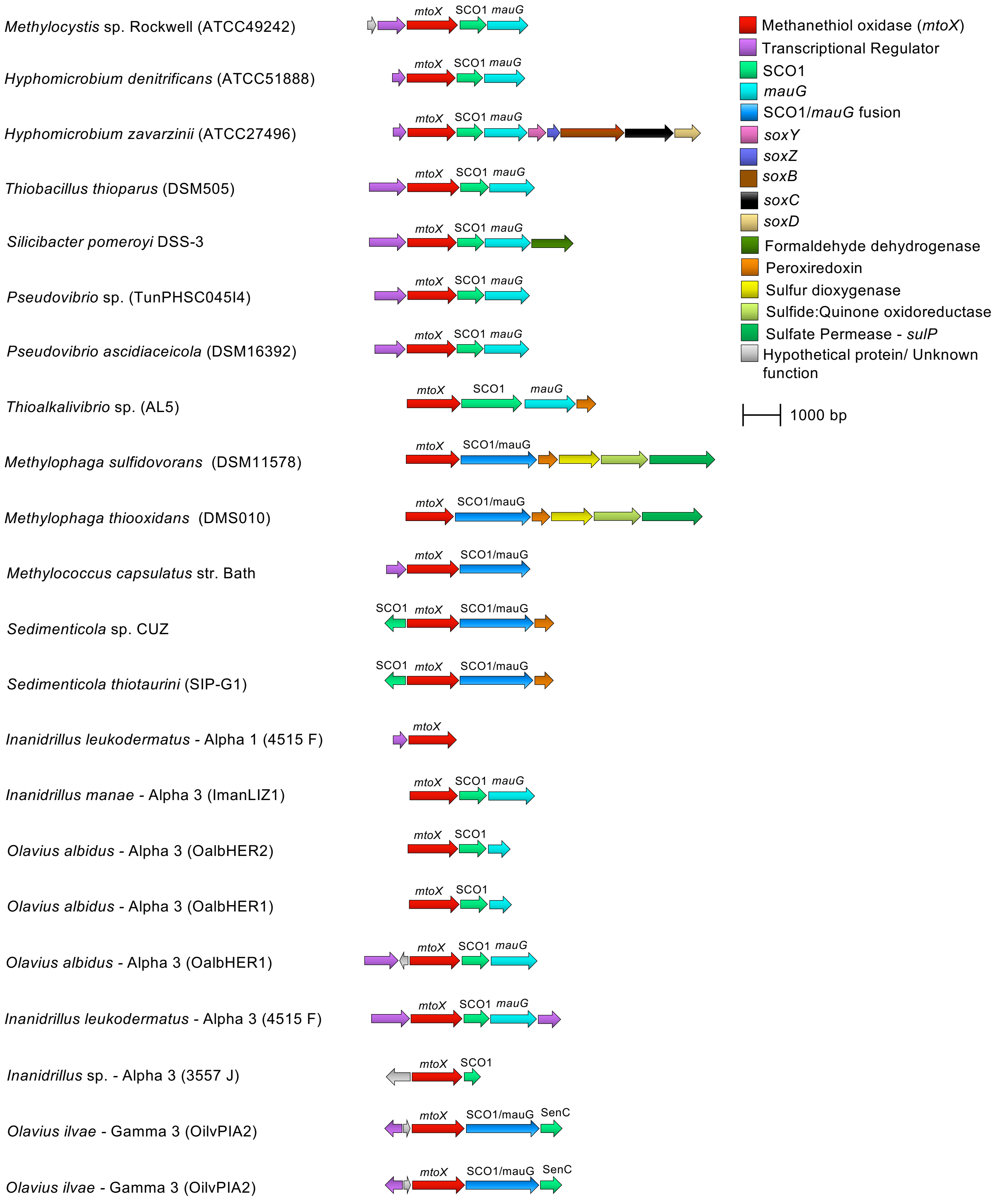

Methylocystis sp. Rockwell (ATCC49242)
Hyphomicrobium denitrificans (ATCC51888)
Hyphomicrobium zavarzinii (ATCC27496)
Thiobacillus thioparus (DSM505)
Silicibacter pomeroyi DSS-3
Pseudovibrio sp. (TunPHSC045I4)
Pseudovibrio ascidiaceicola (DSM16392)
Thioalkalivibrio sp. (AL5)
Methylophaga sulfidovorans (DSM11578)
Methylophaga thiooxidans (DMS010)
Methylococcus capsulatus str. Bath
Sedimenticola sp. CUZ
Sedimenticola thiotaurini (SIP-G1)
Inanidrillus leukodermatus - Alpha 1 (4515 F)
Inanidrillus manae - Alpha 3 (ImanLIZ1)
Olavius albidus - Alpha 3 (OalbHER2)
Olavius albidus - Alpha 3 (OalbHER1)
Olavius albidus - Alpha 3 (OalbHER1)
Inanidrillus leukodermatus - Alpha 3 (4515 F)
Inanidrillus sp. - Alpha 3 (3557 J)
Olavius ilvae - Gamma 3 (OilvPIA2)
Olavius ilvae - Gamma 3 (OilvPIA2)
mtoX
SCO1
mauG
SCO1/mauG
SenC
Methanethiol oxidase (mtoX)
Transcriptional Regulator
SCO1
mauG
SCO1/mauG fusion
soxY
soxZ
soxB
soxC
soxD
Formaldehyde dehydrogenase
Peroxiredoxin
Sulfur dioxygenase
Sulfide:Quinone oxidoreductase
Sulfate Permease - sulP
Hypothetical protein/ Unknown function
1000 bp
