## Supplementary Results for "Chemosynthetic Symbioses as Hidden Hubs of DMSP and Organosulfur Cycling in Marine Sediments"

### Supplementary Results 1

The cleavage pathway of DMSP degradation is represented by the lyase genes (e.g. *dddP* or *dddD*). Across gutless oligochaete symbioses, *dddP* was widespread and phylogenetically diverse, whereas *dddD* was detected only rarely (Fig. 4). In contrast, *dddP* was widespread and phylogenetically diverse, forming several well-supported clusters (Fig. 4 and Fig. S4A). Phylogenetically, gutless-derived *dddD* sequences grouped with characterized lyases from *Ruegeria marina* and *R. pomeroyi*, supporting their role as a functional enzyme converting DMSP to DMS and acryloyl-CoA (Fig. S3A). The *dddP* phylogeny was more complex, comprising one major cluster matching the functionally validated bacterial and fungal DddP clade, which included sequences from alphaproteobacterial and gammaproteobacterial (Gamma3) symbionts (Fig. S4A), and additional clusters of alphaproteobacterial symbionts branching outside this clade but clearly distinct from M24 peptidase outgroups.

Gene neighbourhood analyses revealed distinct metabolic contexts. In references genomes (e.g. Endozoicomonas, Marinomonas*), dddD* co-localized with *dddB* and *dddC*, consistent with acryloyl-CoA processing toward 3-hydroxypropionyl-CoA or 3-hydroxypropionate. In *Ruegeria* spp. and multiple gutless oligochaetes contigs, however, *dddB*/*dddC* were absent within the examined windows (Fig. S3B), suggesting that acryloyl-CoA may be encoded elsewhere in the genome, degraded by general beta-oxidation enzymes, or by metabolic partners within the consortium.

For *dddP, validated references (e.g. Roseobacter denitrificans or Ruegeria pomeroyi)* were adjacent to genes for L-glutamine synthetase, nitrogen regulation, or redox proteins, suggesting integration with nitrogen and redox metabolism. In gutless oligochaete contigs, *dddP* frequently neighboured sarcosine oxidase, dehydrogenase genes, or a TMA methyltransferase-family gene, pointing to links with methylated amine and one-carbon metabolism in the symbioses (Fig. S4B). The high frequency and phylogenetic diversity of *dddP* mirror its prominence in marine metagenomes and indicate a broad capacity for DMSP cleavage among gutless worm symbionts. This widespread *dddP* distribution likely enhances local DMS and acrylate production within host tissues, promoting redox balance and metabolic cross-feeding within the consortium.

**Supplementary Results 2**

The alternative DMSP degradation route, the demethylation pathway, is also well represented, but shows clear taxonomic structuring. Core genes *dmdA* and *dmdB* were detected in 37 and 39 host species, respectively (Fig. 4), though not always together, suggesting partial pathway retention across lineages. Phylogenetic and genomic-context analyses revealed a structured yet cohesive architecture for DMSP demethylation among gutless-oligochaete symbionts. The *dmdA* maximum-likelihood tree resolved three clades: (i) a well-supported group of gutless-derived sequences clustering with canonical references such as *Candidatus* Pelagibacter sp. and *Sulfitobacter* spp., (ii) a large, deeply branching clade dominated by gutless-derived sequences and a single Anaerolineaceae reference; and (iii) a smaller branch of non-DmdA/GcvT-like homologs[1] (Fig. S5A), excluded from downstream analysis due to uncertain function. The placement of many sequences within the canonical DmdA clade and their conserved genomic neighbourhoods confirm their annotation as true DMSP demethylases. While model isolates (e.g., Pelagibacter, Sulfitobacter) encode the full *dmdABCD* operon, gutless symbionts generally exhibit modular rather than operon-like arrangements. In several alphaproteobacterial contigs, *dmdA* is adjacent to *dmdB*, and occasionally to *dmdC* or *acuH*-like hydratase genes. This flexible configuration resembles those seen in cultivated DMSP-degrading bacteria (e.g., *Sulfitobacter geojensis*, *S. guttiformis)*. Additional neighbourhoods place *dmdA* near *acuI*/*acr*-like (acrylyl-CoA reductase) or *acuH*-like (enoyl-CoA hydratase) genes, suggesting links to acrylate detoxification or 3-HP/propionate metabolism. These auxiliary genes likely facilitate recycling of MMPA-CoA and contribute to downstream carbon and sulfur flux within the consortium.

The *dmdB* phylogeny resolved two well-supported clades corresponding to the previously defined B1 and B2 lineages[2] (Fig. S6A). One gutless-derived sequence clustered with Clade 1, whereas the majority belonged to Clade 2. Most *dmdB* homologs originated from alpha- and deltaproteobacterial symbionts, while only a single one was assigned to a gammaproteobacterial symbiont. This contrasts with *dmdA*, which was consistently recovered from alpha- and gammaproteobacterial but not deltaproteobacterial symbionts. Together, these patterns underscore *dmdA* as the key entry-point gene for DMSP demethylation in the alpha- and gammaproteobacterial symbionts of gutless oligochaetes.

By comparison, *dmdB*, which catalyses the CoA-ligation of methylmercaptopropionate (MMPA), shows a broader, taxonomically distinct distribution. Its presence in deltaproteobacterial symbionts suggests these taxa are not directly involved in DMSP demethylation, but may metabolize downstream MMPA. Given that MMPA can also derive from methionine salvage pathways[3], *dmdB* in deltaproteobacterial symbionts may reflect metabolic versatility rather than strict linkage to DMSP turnover.

Overall, the complementary distributions of *dmdA* and *dmdB* indicate a division of labour in DMSP demethylation across the symbiont community. Alphaproteobacterial symbionts encode both genes, enabling complete demethylation and subsequent MMPA metabolism, whereas Gammaproteobacteria appear to initiate DMSP degradation via *dmdA* alone. In contrast, deltaproteobacterial symbionts likely participate in downstream processing of MMPA rather than in the initial demethylation step. Whether the MMPA pool used by these Deltaproteobacteria originates directly from DMSP degradation or from other metabolic sources remains unresolved, but the overall pattern highlights pronounced metabolic complementarity and interdependence within the symbiotic consortium. The lineage-specific presence or absence of *dmdA* and *dmdB* further suggests differential gene retention or loss among symbiont groups (data not shown).

**Supplementary Results 3**

Consistent with its low recovery across metagenomes (Fig. 4), *dmoA* was detected in only a few host species, with a single confidently assigned sequence from *Inanidrillus* sp. (4515C) clustering near canonical *dmoA* sequences (Fig. S7). All other candidates grouped with $FMNH_2$-dependent oxidoreductases distantly related to *bona fide* DmoA, indicating non-specific similarity rather than true homologs. This pattern shows that the DMS monooxygenase pathway is uncommon among gutless oligochaete symbionts.

DMS monooxygenase is a two-component, $FMNH_2$-dependent enzyme system (DmoA/DmoB) that oxidizes DMS to MeSH and formaldehyde[4]. Although homologs occur across diverse proteobacteria and actinobacteria, biochemically confirmed activity is so far limited to *Hyphomicrobium sulfonivorans* and a few additional strains inferred from cell extracts of Hyphomicrobium, Arthrobacter/Pseudarthrobacter, and Thiobacillus species[4-6].

The near-absence of *dmoA* in gutless symbionts suggests that DMS oxidation proceeds predominantly via alternative enzymatic routes, including DMS dehydrogenase (*ddhA*) or trimethylamine monooxygenase (*tmm*), both of which can oxidize DMS to DMSO[7]. A methyltransferase-based route converting DMS to MeSH, proposed in *Methylophaga thiooxydans*[8,9], could represent an additional but unidentified pathway that would not be detectable by our homology-based search, and thus its potential contribution cannot be excluded.

An additional, less-explored route involves the phenol hydroxylase-like system DsoABCDEF[10], whose homologs have been suggested to oxidize DMS to DMSO in certain heterotrophic bacteria. Together, these observations indicate that DMS oxidation in gutless oligochaete symbionts likely relies on DMS dehydrogenase, Tmm, or potentially DsoABCDEF-type hydroxylases, rather than on the classical DmoAB system.

Among the enzymes capable of oxidizing DMS to DMSO, DMS dehydrogenase (*ddhA*) was nearly absent, whereas *tmm* was highly prevalent across gutless oligochaete symbionts (Fig. 4). Only a single *ddhA* sequence, from *Inanidrillus manae*, clustered within the bona fide DdhA clade, while all other candidates grouped within the broader DMSO-reductase family (Fig. S8). *DdhA* encodes the catalytic molybdo-bis(MGD) subunit of DMS dehydrogenase, a periplasmic, multi-subunit enzyme (operon *ddhABDC*) that oxidizes DMS to DMSO (McDevitt, 2002). This enzyme has been functionally characterized in *Rhodovulum sulfidophilum* but occurs in relatively few organisms, consistent with its rarity in our dataset.

In contrast, *tmm* was by far the most frequently recovered DMS-oxidation marker, indicating that this pathway dominates DMS-to-DMSO conversion in gutless symbionts. Tmm is an FAD/NADPH-dependent flavin monooxygenase capable of oxidizing both TMA to trimethylamine-N-oxide (TMAO) and DMS to DMSO, a dual activity experimentally confirmed in *Ruegeria pomeroyi* and other marine bacteria[7].

Phylogenetic analysis revealed a single, well-supported *tmm* clade nested within the broader bacterial FMO family, with gutless oligochaete-derived sequences forming subclades affiliated with Alpha- and Gammaproteobacteria, as well as Spirochaetes (Fig. S9). Multiple *tmm* homologs frequently co-occurred within the same metagenome, indicating contributions from several symbionts and/or paralogous gene copies. The widespread distribution of *tmm* across hosts, comparable to its ~20% representation in marine bacterioplankton genomes, especially among the Rhodobacteraceae (*Roseobacter* group) and SAR11 lineages[11], suggests that gutless symbionts retain analogous metabolic roles. Within these associations, Tmm likely provides both a DMS to DMSO route and a mechanism for TMA oxidation, thereby linking sulfur and nitrogen metabolism.

Flavin monooxygenases (FMOs) are widespread among eukaryotes, including *Caenorhabditis elegans*, where they catalyze the oxygenation of diverse compounds. Given their broad substrate range, these enzymes could in principle perform reactions analogous to bacterial trimethylamine monooxygenase (Tmm). However, their functional equivalence to bacterial Tmm remains unresolved. For example, *C. elegans* expresses at least five FMO paralogs involved in xenobiotic and endogenous amine metabolism[12,13], but none have been biochemically confirmed to oxidize DMS or TMA with the substrate specificity or efficiency characteristic of bacterial Tmm[7,14]. It therefore remains to be established whether eukaryotic FMOs can catalyse comparable reactions, or whether DMS and TMA oxidation in these symbioses is restricted to bacterial enzymes.

**Supplementary Results 4**

DmsABC is a periplasm-facing, membrane-bound molybdoenzyme complex that reduces DMSO to DMS under anoxic or suboxic conditions. It differs from the monomeric DorA/TorA-type reductases and consists of the catalytic subunit DmsA, a multi-[Fe-S] subunit (DmsB), and a membrane anchor (DmsC) that together mediate terminal electron transfer to DMSO during anaerobic respiration[15-17]. The gene *dmsA* was widely represented across hosts metagenomes (Fig. 4), indicating a broad potential for DMSO respiration within gutless oligochaete symbionts. In the maximum-likelihood phylogeny (Fig. S10A), gutless-derived sequences formed well-supported clades within the DMSO reductase enzyme family, mainly assigned to Deltaproteobacteria, with additional sequences from Gamma- and Alphaproteobacteria. Gene-neighbourhood analysis consistently recovered *dmsA* colocalized with *dmsB* and *dmsC*, confirming recovery of the complete *dmsABC* operon in multiple symbionts (Fig. S10B).

Unlike the trimodular DmsABC complex, TorA and DorA are monomeric periplasmic molybdoenzymes that reduce TMAO and/or DMSO. *torA*/*dorA* homologs were frequently detected across hosts metagenomes (Fig. 4), though the number of homologs varied among species. Phylogenetic analyses resolved three major clades within the TorA/DorA lineage (Fig. S11). One clade contained canonical reference enzymes from *Shewanella* spp. (TorA) and *Rhodobacter* spp. (DorA), together with several symbiont sequences. The remaining two sister clades comprised most gutless oligochaete-derived sequences that lack close reference homologs, suggesting the evolution of divergent TorA/DorA-like lineages in these symbioses. Gene-neighbourhood analyses revealed partial conservation of canonical regulatory modules. In several contigs *torA* was situated next to *torR*; one deltaproteobacterial symbiont encoded *torC*, and an alphaproteobacterial representative carried *torR*, *torS* and *torT*. These associations indicate preservation of key regulatory and electron-transfer components, although complete operons were not observed. The alphaproteobacteria-dominant distribution of TorA/DorA versus the deltaproteobacteria-dominated distribution of DmsABC suggests lineage-specific respiratory strategies for handling N-oxides and DMSO.

In addition to enzymatic formation from DMS via Tmm, DMSO can also originate from abiotic oxidation of DMS or enzymatic cleavage of dimethylsulfoxoniopropionate (DMSOP), an oxidised form of DMSP. While DMSOP concentrations in pelagic waters are typically lower than those of DMSP[18], initial measurements in marine sediments suggest that DMSOP levels may be comparable to or even exceed those of DMSP. Although DMSO concentrations have not yet been measured in sediments inhabited by gutless oligochaetes, data from intertidal mudflat and saltmarsh sediments show surface concentrations averaging ~11 nmol $g^{-1}$ and reaching >100 nmol $g^{-1}$ [19]. These findings imply that, in addition to DMSO generated by microbial DMSP and

DMS metabolism, substantial environmental DMSO is potentially available as an external electron acceptor. The presence of *tmm* and *dmsA* in gutless oligochaete metagenomes supports the potential for a local DMS-DMSO redox loop within the holobiont. Such cycling would provide symbionts with a flexible electron-acceptor system, stabilize redox balance, and modulate DMS fluxes under fluctuating sediment conditions.

Beyond DMS and DMSO cycling, DMSP and DMS oxidation also generate methanethiol (MeSH). To investigate its fate in the symbioses, we next examined the distribution of methanethiol oxidase (MtoX).

**Supplementary Results 5**

MtoX is a copper-dependent periplasmic oxidase of the SBP56 family that converts methanethiol (MeSH) to formaldehyde, hydrogen peroxide, and a reduced sulfur product[20]. While earlier work identified this product as $H_2S$, recent biochemical evidence indicates that sulfane sulfur ($S^o$) is the main product, which can subsequently be oxidized to sulfate or incorporated into other sulfur species[21]. Methanethiol oxidase (*mtoX*) was moderately common across gutless oligochaete hosts (Fig. 4), detected in 19 species. Phylogenetic analysis resolved two major clades corresponding to Alphaproteobacteria and Gammaproteobacteria (Fig. S12A). Alpha1 and Alpha3 symbionts (*Olavius albidus*, *Inanidrillus manae*, *I. leukodermatus*) formed a sister clade to free-living Alphaproteobacteria, which typically encode *mtoX* adjacent to separate *SCO1* and *mauG* genes (Fig. S12B)[20]. In contrast, Gamma3 symbionts clustered with free-living Gammaproteobacteria, where *mtoX* is often, though not universally, linked to a fused *mauG-SCO1* gene (e.g., *Thioalkalivibrio* retains the separated configuration). The consistent co-occurrence of *mtoX* with its presumed maturation partners *mauG* and *SCO1* (sometimes fused) indicates that canonical biogenesis and copper delivery systems are preserved in gutless symbionts. The formation of a mainly gutless-specific phylogenetic branch suggests either host-associated diversification of this enzyme or underrepresentation of similar environmental lineages in current databases.

In addition to bacterial *mtoX*, our expanded phylogeny (Fig. S12A) also included sequences from animal genomes, revealing that the *O. algarvensis* host encodes a SELENBP1-like methanethiol oxidase homolog. This host sequence grouped within the eukaryotic SELENBP1 clade together with human[22] and *Caenorhabditis elegans*[23] proteins, distinct from bacterial *mtoX*. Its phylogenetic placement and conserved active-site residues indicate that the host likely possesses methanethiol-oxidizing capacity. This provides independent genomic support for a potential host contribution to MeSH detoxification, consistent with the metabolic handoff proposed for the holobiont. In summary, *mtoX* is a moderately abundant yet phylogenetically cohesive marker of methanethiol oxidation in gutless oligochaete symbionts. Its conserved genomic context and retention of maturation partners support a central role in MeSH detoxification and redox balance within the consortium, potentially via sulfane-sulfur production that can re-enter the internal sulfur cycle[21].
